# Representing linguistic communicative goals in the premotor cortex

**DOI:** 10.1101/2022.05.26.493580

**Authors:** Wenshuo Chang, Lihui Wang, Ruolin Yang, Xingchao Wang, Zhixian Gao, Xiaolin Zhou

**Author notes:** Address correspondence to Dr. Lihui Wang, Dr. Xingchao Wang, or Prof. Xiaolin Zhou.

## Abstract

Linguistic communication is often regarded as an action conveying the speaker’s communicative goal to the addressee. With both correlational (an fMRI study) and causal (a lesion study) evidence, we demonstrated that communicative goals are represented in human premotor cortex. Participants read scripts each containing a sentence said by the speaker with a goal of either a promise, a request, or a reply. The fMRI results showed that the premotor cortex represented more information on communicative goals than the perisylvian language regions. The lesion study results showed that, relative to healthy controls, the understanding of communicative goals was impaired in patients with lesions in the premotor cortex, whereas no reliable difference between the healthy controls and lesion controls. These findings convergently suggest that the premotor cortex is crucial for representing the goals conveyed by language, supporting the theoretical view that linguistic communication can be seen as a goal-directed action.

## Introduction

Linguistic communication usually engages two interlocutors, a speaker and an addressee (Brennan et al., 2010; Russell, 1950; Tylén et al., 2010). Linguistic theories, such as Sprachspiel (Wittgenstein, 1953) and speech acts theory (Austin, 1975; Searle, 1969; 1985), propose that language use is a communicative action conveying the speaker’s goal, which expresses what is intended to be achieved, to the addressee. If linguistic communication is analogized as electric circuit, the speaker’s utterance is the wire connecting the interlocutors’ minds, and the goal is the current goes through the wire to transfer information. To interpret the transferred information, understanding the speaker’s goal is an essential step for the addressee who seeks to respond accordingly (Levinson, 2016). In other words, correctly understanding the speaker’s goal is critical for a successful communication.

Depending on the speaker’s goals (Searle, 1969; 1985), linguistic communications can be categorized into commissives, directives, and assertives. By commissives, such as to promise or to assure, the speaker shows his/her commitment to conduct a task. By directives, such as to request or to order, the speaker obliges the addressee to conduct a task. While both commissives and directives involve conducts of tasks which serve the intentions of the interlocutors, assertives, such as replying to a query or stating a fact, involve the description of objective situations which can be irrelevant to the interlocutors’ intentions. These categories can be differentiated by the interlocutors’ attitudes toward the tasks, such as their willingness and their evaluations on the cost-benefit of accomplishing the task (Pérez Hernández, 2001; Searle and Vanderveken, 1985). For example, by saying “I will write a recommendation letter for you” as a promise, the speaker conveys a goal that can benefit the addressee, and hence the addressee would have willingness to have the task accomplished by the speaker. By saying “please write a recommendation letter for me” as a request, the speaker conveys a goal that can benefit the speaker, and hence the speaker would have willingness to have the task accomplished. By saying “I wrote a recommendation letter for my student” as a reply to the question “what did you do this morning?” the interlocutors have no clear attitude towards the task in the reply.

While linguistic communications are regarded as goal-directed actions in linguistic theories, it is barely known how these communicative goals could be represented in the brain. One straightforward prediction is that communicative goals are represented in brain areas that subserve action programing or preparation. Consistent with this prediction, the co-evolution of human’s linguistic ability and motor skills (e.g., tool use) has been highlighted from neurophysiological, neurocognitive, and anthropological perspectives (Arbib, 2011; Pulvermüller, 2018; Rizzolatti and Arbib, 1998; Stout and Chaminade, 2012; Thibault et al., 2021). As a demonstration, linguistic communications between tutors and learners can improve the efficiency of learning to make Palaeolithic tools (Morgan et al., 2015), and the activation in the premotor region of the human brain increases with the evolutionary progress of Palaeolithic tool-making skills (Stout et al., 2008). Moreover, contributions of the premotor region to communications through language or language-like manners are revealed not only in humans (Dreyer and Pulvermüller, 2018; Egorova et al., 2016; Hauk et al., 2004; Wilson et al., 2004), but also in species including avian (Thompson et al., 2011), Cercopithecinae (Gil-da-Costa et al., 2006), and Pan troglodytes (Bianchi et al., 2016).

For humans, the premotor cortex, consisting of the lateral premotor cortex (LPMC) and medial premotor cortex (MPMC) (Mayka et al., 2006), are broadly involved in action-related processes, such as action execution (Aziz-Zadeh et al., 2006), planning (Gallivan et al., 2013), observation (Aziz-Zadeh *et al*., 2006), imitation (Aziz-Zadeh *et al*., 2006), and imagery (Pilgramm et al., 2016). Importantly, the premotor cortex is also found to play a crucial role in human language processing (Arbib, 2011; 2016; Gallese, 2008; Gallese and Lakoff, 2005; Hertrich et al., 2016; Pulvermüller, 2005; 2018; Pulvermüller and Fadiga, 2010). The premotor cortex is involved in speech production and perception which engage explicit motor programing of articulator organs (Wilson *et al*., 2004), as well as in language comprehension without such explicit programing (Feng et al., 2017; Feng et al., 2021; Hauk *et al*., 2004; Postle et al., 2008; van Ackeren et al., 2012). For example, the comprehension of written action verbs involves the activation of the premotor cortex (Hauk *et al*., 2004), and the processing of action semantics is interfered by transcranial magnetic stimulation (TMS) over the premotor cortex (Courson et al., 2017; Willems et al., 2011).

Moreover, the premotor cortex is not only involved in the language processing that is directly related to action semantics, but also involved in linguistic communications that are not literally related to actions (Feng *et al*., 2021; Shibata et al., 2011; van Ackeren *et al*., 2012). Relative to hearing statements of objective situations (e.g., ‘It is hot here’ with a picture of a desert), hearing indirect requests (e.g., ‘It is hot here’ with a picture of a closed window) elicits greater activations in the left LPMC and left MPMC (van Ackeren *et al*., 2012). Similarly, the indirect reply (e.g., saying “it’s hard to give a good presentation”) to the addressee’s question (e.g., “what did you think of my presentation?”) elicits increased activation in the medial frontal cortex extending to the MPMC as compared with a literal reply (Feng *et al*., 2021; Shibata *et al*., 2011). Relative to hearing prosodies conveying unambiguous communicative goals, hearing prosodies conveying ambiguous goals engenders stronger activation in the MPMC (Hellbernd and Sammler, 2018). Moreover, the premotor cortex is responsive not only to the ambiguity of the communicative goals but also to the types of communicative goals (Egorova *et al*., 2016). Relative to a verbal assertive, a verbal request elicits increased activations in the bilateral premotor cortex as well as in the left inferior frontal gyrus and temporal regions.

Given the usual concurrent involvement of the premotor cortex and the left lateralized perisylvian language regions (Egorova *et al*., 2016; Feng *et al*., 2017; Feng *et al*., 2021; Shibata *et al*., 2011), the latter of which are broadly involved in the processing of semantic and syntactic information (Friederici, 2011; Friederici et al., 2017; Hagoort, 2017), it is unknown if the premotor cortex functions more or less profoundly in representing communicative goals than the perisylvian regions. The present study aims to test to what extent the premotor cortex plays a critical role in representing different communicative goals when linguistic communications are understood. To this end, we first conducted an fMRI study in which participants were instructed to read scripts of linguistic communications, each of which contained a specific communicative goal of the speaker. We expected that the different communicative goals conveyed in the scripts can be decoded by the multivariate brain activation patterns. Considering the role of the premotor cortex in representing action-related information, we predicted further that the decoding of communicative goals is more pronounced in the premotor cortex than in the perisylvian regions. Moreover, we predicted that the representation of communicative goals in the premotor cortex is correlated with the interlocutors’ attitudes which are closely related to communicative goals. We also conducted a lesion study on patients with brain lesions to assess the causal role of the premotor cortex in representing communicative goals. We predicted that the understanding of communicative goals would be impaired in the patient group with lesions in the premotor cortex, relative to the patient group with lesions in other brain areas and to the healthy control group.

## Results

### Behavioral results of the fMRI study

We created 80 Chinese scripts, each of which began with a context introducing two interlocutors, followed by a pre-critical sentence specifying the identity of the interlocutors (i.e., the speaker and the addressee). The script ended with a critical sentence which was the speaker’s utterance to the addressee. As shown in Table 1, depending on the context, the critical sentences conveyed different communicative goals: 1) a promise (*Promis*e condition); 2) a reply to the addressee’s query (*Reply-1* condition), serving as a control to the *Promise* condition; 3) a request (*Request* condition); 4) a reply to the addressee’s query (*Reply-2* condition), serving as a control to the *Request* condition. The validity of these scripts in reflecting communicative goals was confirmed by pilot evaluative results (*Supplemental Information*). In the pilot evaluation, for each script, participants rated the extent to which communicative goals could be predicted by the features revealed by Pérez Hernández’s corpus study (2001): speaker’s will, speaker’s cost-benefit, speaker’s pleasure, addressee’s will, addressee’s cost-benefit, addressee’s pleasure, the performer’s capability, relative power and social distance between the interlocutors, and the mitigation of the critical sentence. An exploratory factor analysis (EFA) revealed three accounting factors for these features: the speaker’s attitudes, covering speaker’s will, cost-benefit, and pleasure; the addressee’s attitudes, covering addressee’s will, cost-benefit, and pleasure; and contextual information, covering relative power, social distance, and the mitigation (see *Supplemental Information* for the statistics). The feature of the performer’s capability had loadings lower than 0.3 on any of the three accounting factors and hence was not considered as explainable by either factor.

**Table 1.** English translation of the scenarios in the fMRI study, with original Chinese version of critical sentences.

| Communicative goal | Context | Pre-critical sentence | Critical sentence |
| --- | --- | --- | --- |
| *Promise* | The sales department conducted a survey, and Xiaoli was assigned to analyze the surveyed data this week. But Xiaoli was too busy to analyze it because he had the other assignment recently. His colleague Xiaowu had more spare time this week. And they were communicating with each other. | Then Xiaowu said to Xiaoli: | “I will analyze the survey data this week.”<br>“这周我负责分析问卷数据。” |
| *Reply-1* | The sales department conducted a survey, and Xiaowu was assigned to analyze the surveyed data this week. His colleague Xiaoli was assigned to record a working memo, Xiaoli wanted to know about Xiaowu’s job assignment. And they were communicating with each other. | Then Xiaowu said to Xiaoli: | “I will analyze the survey data this week.”<br>“这周我负责分析问卷数据。” |
| *Request* | The sales department conducted a survey, and Xiaowu was assigned to analyze the surveyed data this week. But Xiaowu was too busy to analyze the surveyed data because he had the other assignment recently. His colleague Xiaoli had more spare time this week. And they were communicating with each other. | Then Xiaowu said to Xiaoli: | “You will analyze the survey data this week.”<br>“这周你负责分析问卷数据。” |
| *Reply-2* | The sales department conducted a survey, and Xiaoli was assigned to analyze the surveyed data this week. His colleague Xiaowu was assigned to record a working memo. Then, Xiaoli wanted to confirm his own job assignment. They were communicating with each other. | Then Xiaowu said to Xiaoli: | “You will analyze the survey data this week.”<br>“这周你负责分析问卷数据。” |

The reliability and the validity of these scripts were further confirmed by the same pattern of rating results obtained from the fMRI participants after the fMRI scanning. Bayesian logistic mixed models were applied on the post-scanning ratings to estimate model coefficients of the features in predicting the pair-wise communicative goals, “*Promise* vs. *Reply-1*” and “*Request* vs. *Reply-2*”, respectively (Figure 1b and *Table S3*). Results showed that, for “*Promise* vs. *Reply-1*”, the ratings of addressee’s will, cost-benefit, and pleasure were higher for *Promise* than for *Reply-1* while the ratings of speaker’s cost-benefit and social distance were lower for *Promise* than for *Reply-1*; for “*Request* vs. *Reply-2”*, the ratings of speaker’s will, cost-benefit, and pleasure and relative power were higher for *Request* than for *Reply-2* while the ratings of addressee’s will, cost-benefit, and pleasure and the mitigation were lower for *Request* than for *Reply-2*. A further confirmatory factor analysis (CFA) on the post-scanning ratings was conducted to confirm the appropriateness of the three-factor model obtained by the above EFA and to show a good fit of the three-factor model (Figure 1c; *CFI* = 0.92, TLI = 0.88, *RMSEA* = 0.11).

**Figure 1.**
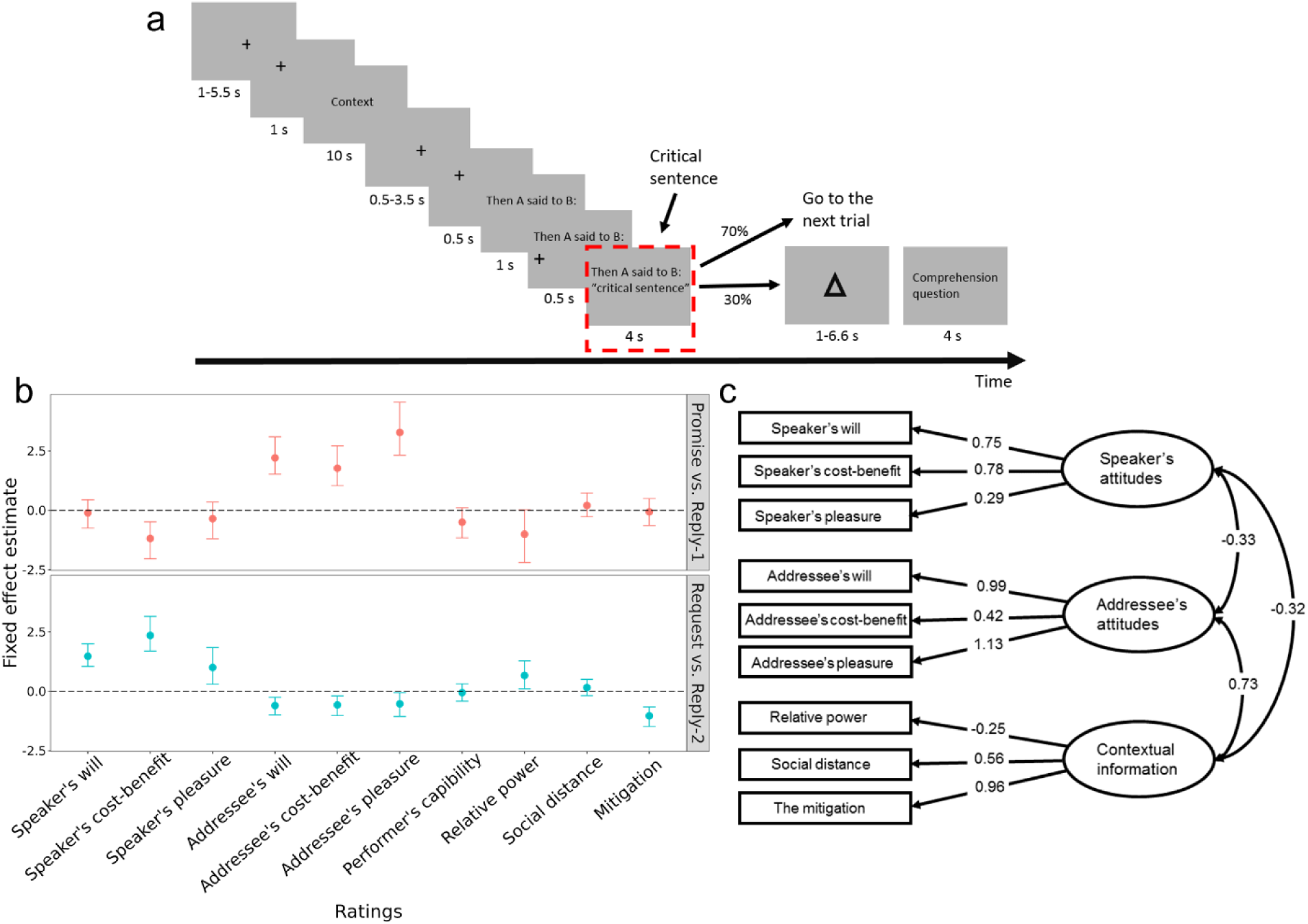
Experimental procedure and behavioral results of the fMRI study. **a,** In each trial, the context, the pre-critical sentence, and the critical sentence were presented sequentially in written form. The critical sentence is enclosed by the dashed red rectangle (not shown in the actual experiment). In 24 catch trials (30% of all trials), participants were instructed to respond to a comprehension question. **b,** Results of Bayesian logistic mixed modelling. The posterior estimate of the fixed effect (vertical axis) for each feature (horizontal axis) in the “*Promise* vs. *Reply-1*” model (upper panel, red) and the “*Request* vs. *Reply-2*” model (lower panel, turquoise) are illustrated. The solid dots represent mean posterior estimates, the error bars represent 95% credible intervals (*CrI*s). A 95% *CrI* excluding 0 indicates a statistically significant predictability of the corresponding feature. **c,** The three-factor model of the CFA. The ellipses represent the accounting factors and the rectangles represent the features. The correlations between the accounting factors and the loadings of the accounting factors on the features are embedded in the arrows.

### Multivariate pattern classifications (MVPCs) of fMRI data

In the fMRI experiment, each participant read the written scripts during the fMRI scanning (Figure 1a). We asked if the critical sentences that conveyed different communicative goals induced different patterns of BOLD signals in the premotor cortex that could be discriminated by multivariate pattern classification (MVPC). The univariate analysis did not reveal significant effects related to the aim of this study (*Supplemental Information*). Regions of interest (ROIs) within the perisylvian regions, including the BA44 division of Broca’s area (left BA44), the BA45 division of Broca’s area (left BA45), left middle temporal gyrus (LMTG), and left superior temporal gyrus (LSTG) (Friederici *et al*., 2017), were defined in addition to the premotor cortex. The lateral premotor cortex (LPMC) and medial premotor cortex (MPMC) were separately defined (Mayka *et al*., 2006) and analyzed to avoid the risk of overfitting because the combined lateral and medial parts would contain more voxels than each of the other ROIs beyond an order of magnitude (Figure 2a).

**Figure 2.**
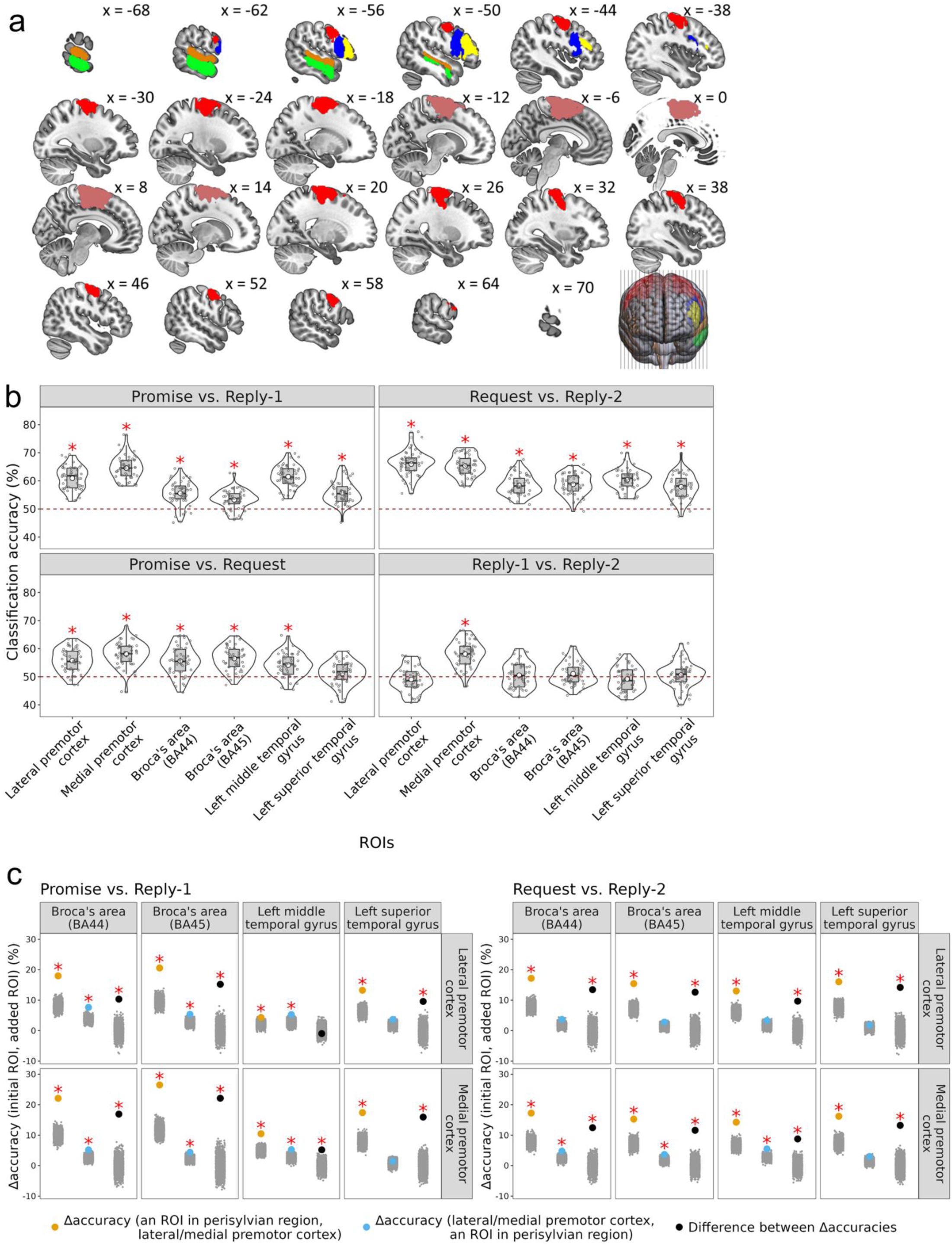
MVPCs of fMRI data. **a,** Five ROIs were defined. Red, lateral premotor cortex; pink, medial premotor cortex; blue, Broca’s area (BA44); yellow, Broca’a area (BA45); green, left middle temporal gyrus; orange, left superior temporal gyrus (x-coordinates based on the MNI system). **b,** Results of ROI-based MVPCs. The classification accuracies (vertical axis) in the ROIs (horizontal axis) for the four pair-wise classifications are illustrated. Left top, *Promise* vs. *Reply-1*; right top, *Request* vs. *Reply-2*; left bottom, *Promise* vs. *Request*; right bottom, *Reply-1* vs. *Reply-2*. The red dashed lines represent the chance-level percentage of binary classification (50%). Red stars represent the statistical significance of permutation tests with Bonferroni correction. **c,** Results of combinatorial MVPCs. Left panel, *Promise* vs. *Reply-1*; right panel, *Request* vs. *Reply-2.* Vertical axes illustrate the improvement in classification accuracy contributed by an added ROI for an initial ROI. Red stars represent statistical significance of permutation tests. Each yellow dot indicates the improvement in classification accuracy contributed by a premotor ROI for a perisylvian ROI. Each blue dot indicates the improvement in classification accuracy contributed by a perisylvian ROI for a premotor ROI. Each black dot indicates the difference between the improvement in classification accuracy contributed by a premotor ROI for a perisylvian ROI and that contributed by a perisylvian ROI for a premotor ROI. The crowded small gray dots indicate data points of null distributions for permutation tests.

As shown in Figure 2b, for “*Promise* vs. *Reply-1*”, classification accuracies were above chance level in all ROIs (all *p*-values < .0005, permutation-based significance testing with Bonferroni corrections for multiple comparisons, Table 2). For “*Request* vs. *Reply-2*”, accuracies were also above chance level in all ROIs (all *p*-values < .0005). For “*Promise* vs. *Request*”, accuracies were above chance level in all ROIs (all *p*-values < .0005) except LSTG. For “*Reply-1* vs. *Reply-2*”, the accuracy was above chance only in MPMC (*p*-value < .0005).

**Table 2.** Results of MVPCs.

| ROI-based MVPCs |  |  |  |  |  |  |  |  |
| --- | --- | --- | --- | --- | --- | --- | --- | --- |
| Pair-wise classification |  |  |  |  |  |  |  |  |
| ROI (n volxels) | Index |  | *Promise vs. Reply-1* | *Request vs. Reply-2* | *Promise vs. Request* | *Reply-1 vs. Reply-2* |  |  |
| LPMC (5507) | *Accuracy (%)* |  | **60** | **67** | **55** | 50 |  |  |
|  | *p* |  | **< 0.0005** | **< 0.0005** | **< 0.0005** | 0.534 |  |  |
| MPMC (5227) | *Accuracy (%)* |  | **64** | **67** | **58** | **58** |  |  |
|  | *p* |  | **< 0.0005** | **< 0.0005** | **< 0.0005** | **< 0.0005** |  |  |
| Left BA44 (1443) | *Accuracy (%)* |  | **56** | **58** | **55** | 51 |  |  |
|  | *p* |  | **< 0.0005** | **< 0.0005** | **< 0.0005** | 0.128 |  |  |
| Left BA45 (979) | *Accuracy (%)* |  | **53** | **58** | **58** | 49 |  |  |
|  | *p* |  | **< 0.0005** | **< 0.0005** | **< 0.0005** | 0.827 |  |  |
| LMTG (1760) | *Accuracy (%)* |  | **61** | **60** | **55** | 51 |  |  |
|  | *p* |  | **< 0.0005** | **< 0.0005** | **< 0.0005** | 0.078 |  |  |
| LSTG (1147) | *Accuracy (%)* |  | **56** | **57** | 51 | 51 |  |  |
|  | *p* |  | **< 0.0005** | **< 0.0005** | 0.018 | 0.028 |  |  |
| Combinatorial MVPCs of Δaccuracy (*initial ROI, added ROI*) | | | | | | | | |
| Pair-wise classification | Premotor ROI | Perisylvian ROI | Δaccuracy (*perisylvian ROI, premotor ROI*) | | Δaccuracy (*premotor ROI, perisylvian ROI*) | | Difference between Δaccuracies | |
| | | | Δaccuracy (%) | *p* | Δaccuracy (%) | *p* | Δaccuracy (%) | *p* |
| *Promise*<br>vs.<br>*Reply-1* | LPMC | Left BA44 | **18** | **< 0.0005** | **8** | **< 0.0005** | **10** | **< 0.0005** |
|  |  | Left BA45 | **21** | **< 0.0005** | **5** | **< 0.0005** | **16** | **< 0.0005** |
|  |  | LMTG | **4** | **0.0015** | **5** | **< 0.0005** | -1 | 0.5 |
|  |  | LSTG | **13** | **< 0.0005** | 4 | 0.0045 | **9** | **< 0.0005** |
|  | MPMC | Left BA44 | **22** | **< 0.0005** | **5** | **< 0.0005** | **17** | **< 0.0005** |
|  |  | Left BA45 | **26** | **< 0.0005** | **4** | **0.001** | **22** | **< 0.0005** |
|  |  | LMTG | **10** | **< 0.0005** | **5** | **< 0.0005** | **5** | **< 0.0005** |
|  |  | LSTG | **17** | **< 0.0005** | 1 | 0.1565 | **16** | **< 0.0005** |
| *Request*<br>vs.<br>*Reply-2* | LPMC | Left BA44 | **17** | **< 0.0005** | 4 | 0.002 | **12** | **< 0.0005** |
|  |  | Left BA45 | **15** | **< 0.0005** | 3 | 0.0115 | **12** | **< 0.0005** |
|  |  | LMTG | **13** | **< 0.0005** | 3 | 0.0055 | **10** | **< 0.0005** |
|  |  | LSTG | **16** | **< 0.0005** | 2 | 0.0725 | **14** | **< 0.0005** |
|  | MPMC | Left BA44 | **17** | **< 0.0005** | **5** | **< 0.0005** | **12** | **< 0.0005** |
|  |  | Left BA45 | **15** | **< 0.0005** | **4** | **0.0015** | **11** | **< 0.0005** |
|  |  | LMTG | **14** | **< 0.0005** | **5** | **< 0.0005** | **9** | **< 0.0005** |
|  |  | LSTG | **16** | **< 0.0005** | 3 | 0.01 | **13** | **< 0.0005** |

Texts in bold indicate statistical significance for the permutation-based significance testing with Bonferroni correction.
For ROI-based MVPCs, accuracies were tested with Bonferroni corrected significance threshold of $p < 0.002$ . For combinatorial MVPCs, statistical significances for the permutation tests at the first step and the second step were determined by Bonferroni corrected thresholds of $p < 0.0016$ and $p < 0.003$ respectively. LPMC, lateral premotor cortex; MPMC, medial premotor cortex; Left BA44, Broca's area (BA44); Left BA45, Broca's area (BA45); LMTG, Left middle temporal gyrus; LSTG, Left superior temporal gyrus.

To examine whether the premotor cortex represented more information on communicative goals than the perisylvian regions, we conducted combinatorial MVPCs (Carter et al., 2012; Clithero et al., 2009). This was implemented by quantifying the extent to which an added ROI improved the classification accuracy of an initial ROI. At the first step, using one of the perisylvian ROIs as the initial ROI and either LPMC or MPMC as the added ROI, we showed that the accuracies in classifying “*Promise* vs*. Reply-1*” (Figure 2c left panel and Table 2) were significantly improved by adding LPMC, or by adding MPMC. The accuracies in classifying “*Request* vs. *Reply-2*” (Figure 2c right panel and Table 2) were also significantly improved by adding LPMC or MPMC.

Using LPMC or MPMC as the initial ROI and one of the perisylvian ROIs as the added ROI, we showed that, in classifying “*Promise* vs. *Reply-1*” (Figure 2c left panel and Table 2), adding either left BA44, left BA45, or LMTG significantly improved the accuracies in LPMC or MPMC. In classifying “*Request vs. Reply-2*” (Figure 2c right panel and Table 2), adding either left BA44, left BA45, or LMTG significantly improved the accuracies in MPMC, but not in LPMC.

At the second step, we compared the improvements in classification contributed by the premotor cortex with the improvements contributed by the perisylvian regions. For “*Promise vs. Reply-1*” (Figure 2c left panel and Table 2), the results showed that the improvement contributed by LPMC for either of the perisylvian ROIs (except LMTG) was larger than the improvement contributed by either of the perisylvian ROIs for LPMC (left BA44: 18% vs. 8%; left BA45: 21% vs. 5%; LSTG: 13% vs. 4%, all *p*-values < 0.0005); the improvement contributed by MPMC for either of the perisylvian ROIs was larger than the improvement contributed by either of the perisylvian ROIs for MPMC (left BA44: 22% vs. 5%; left BA45: 26% vs. 4%; LMTG: 10% vs. 5%; LSTG: 17% vs. 1%, all *p*-values < 0.0005).

For “*Request vs. Reply-2*” (Figure 2c right panel and Table 2), the same pattern of results was observed on the improvements contributed by LPMC for all the perisylvian ROIs (left BA44: 17% vs. 4%; left BA45: 15% vs. 3%; LMTG: 13% vs. 3%; LSTG: 16% vs. 2%, all *p*-values < 0.0005); and on the improvements contributed by MPMC (left BA44: 17% vs. 5%; left BA45: 15% vs. 4%; LMTG: 14% vs. 5%; LSTG: 16% vs. 3%, all *p*-values < 0.0005).

In addition, as medial prefrontal cortex (MPFC) and temporo-parietal junction (TPJ) were shown to activate in processing linguistic communications in previous studies (e.g., indirect reply, Feng *et al*., 2017; Feng *et al*., 2021; Shibata *et al*., 2011), we compared the amount of information on communicative goals represented in the premotor cortex with the amount of information represented in MPFC and TPJ, using the same methods illustrated above (*Supplemental Information*). The results suggested that, although MPFC and TPJ represent information on communicative goals to a certain extent, the premotor cortex represents more (*Figure S4* and *Table S3*).

Taken together, the MVPC results suggested that while both the premotor cortex and the perisylvian regions contain information on communicative goals, the premotor cortex represented more information relative to the perisylvian regions and other brain areas previously shown to be related to linguistic communications.

### Representational similarity analysis (RSA) results of fMRI data

As shown by the rating results, communicative goals are related to the interlocutor-related features. We thus conducted RSA (Kriegeskorte et al., 2008) to test to what extent the activation pattern in the premotor cortex and in the perisylvian regions could be predicted by the three behavioral accounting factors (speaker’s attitudes, addressee’s attitudes, and contextual information), and vice versa.

First, for the brain representational dissimilarity matrix (RDM) of each ROI and each of the two pair-wise predictions, “*Promise* vs. *Reply-1*” and “*Request* vs. *Reply-2*”, a representational similarity (RS) encoding model was fitted with the brain RDM being included as the response variable and the three behavioral RDMs as the predictors (Figure 3a and Figure 3b). For “*Promise* vs. *Reply-1*” (Figure 3c upper panel and Table 3), each brain RDM was significantly predicted by Addressee RDM, but not by Speaker RDM or Context RDM. For “*Request vs. Reply-2*” (Figure 3c lower panel and Table 3), brain RDMs of LPMC, MPMC, left BA44, LMTG, and LSTG were significantly predicted by Speaker RDM, but not by Addressee RDM or Context RDM. However, no significant effect was observed for left BA45 RDM.

**Figure 3.**
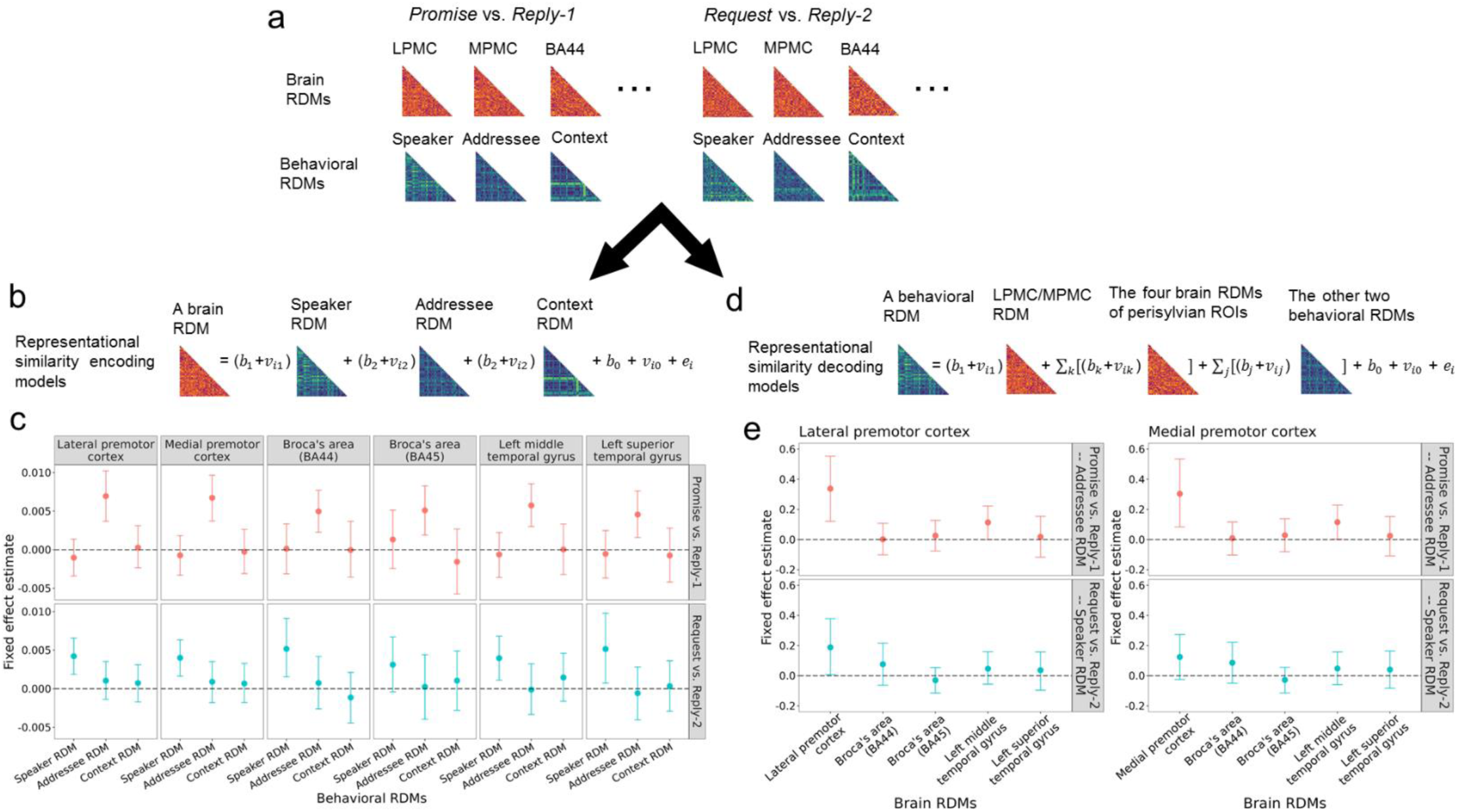
RSAs of fMRI data. **a,** The brain RDMs and the behavioral RDMs for the two pair-wise predictions, “*Promise* vs. *Reply-1*” and “*Request* vs. *Reply-2*”. **b,** The equation of the representational similarity encoding model, which includes a brain RDM as response variable and the three behavioral RDMs as predictors (see *Methods* for details). **c,** Results of representational similarity encoding models. **d,** The equation of the representational similarity decoding model, which includes a behavioral RDM as response variable, five brain RDMs as predictors, and the other two behavioral RDMs as covariates (see *Methods* for details). **e,** Left panel, results of representational similarity decoding models with the LPMC RDM and the RDMs of the prerisylvian ROIs as predictors; right panel, results of representational similarity decoding models with the MPMC RDM and the RDMs of the perisylvian ROIs as predictors. For **a**, **b,** and **d**, the lower-triangular RDMs from one participant are shown as examples (only for illustrative purpose). For **c** and **e**, the posterior estimates (vertical axis) of the “*Promise* vs. *Reply-1*” models (upper panel, red) and the “*Request* vs. *Reply-2*” models (lower panel, turquoise) are illustrated. The solid dots represent mean posterior estimates, the error bars represent Bonferroni corrected *CrI*s (99.86% *CrI* for the encoding models and 99.92% *CrI* for the decoding models). The dashed gray lines indicate 0 for fixed effect estimates. An effect was determined as significant when the Bonferroni-corrected *CrI* excluded 0.

**Table 3.**
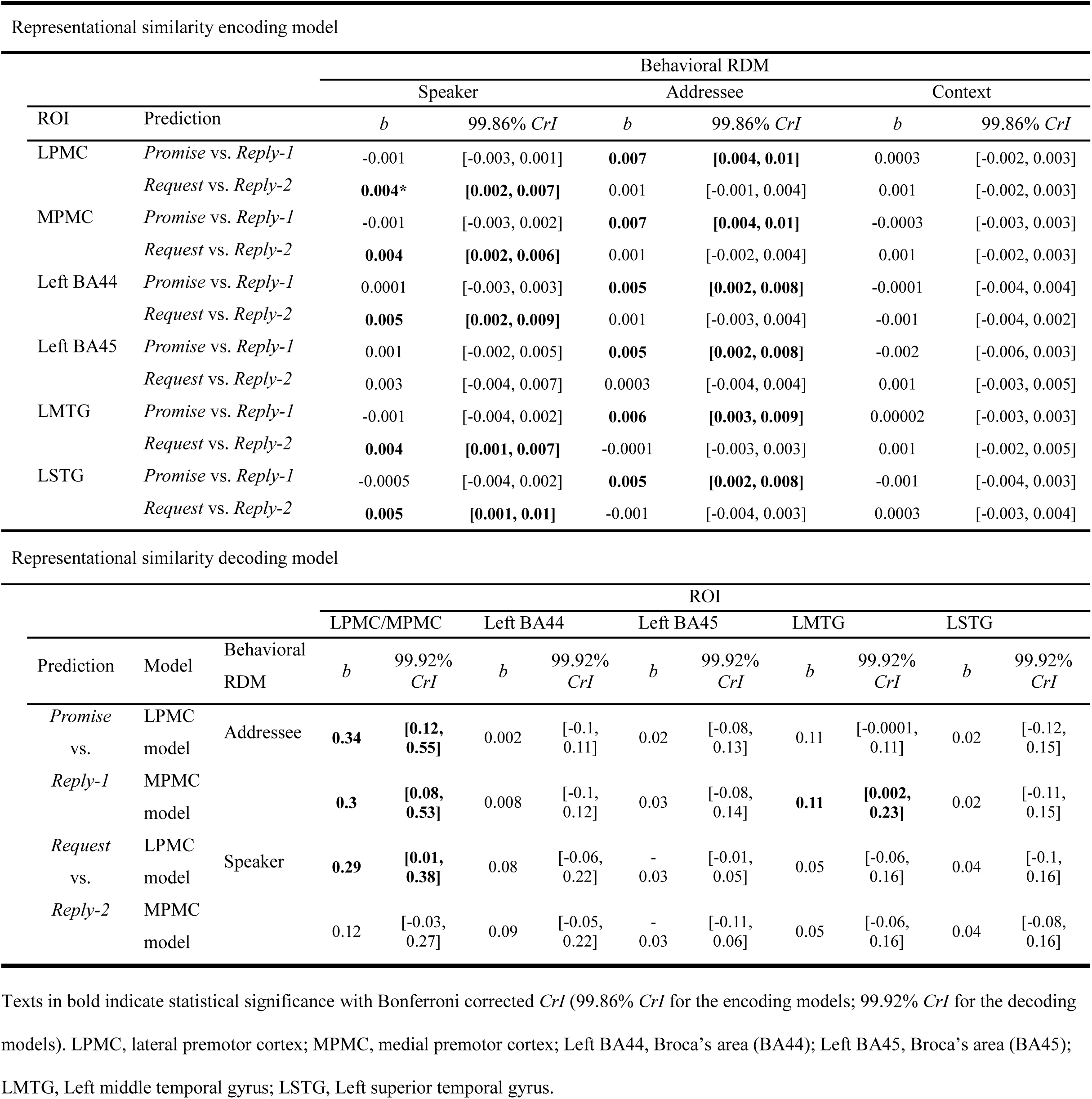
Results of RSAs.

Second, for each behavioral RDM and each pair-wise prediction, an RS decoding model was fitted with a behavioral RDM (Speaker RDM, Addressee RDM, or Context RDM) as the response variable, and the brain RDMs of LPMC/MPMC and the perisylvian ROIs as the predictors. The other two behavioral RDMs were included as covariates (Figure 3d). Note that LPMC and MPMC were modeled separately to avoid co-linearity. Hence, there were LPMC models and MPMC models.

For “*Promise* vs. *Reply-1*” (Figure 3e left panel and Table 3), Addressee RDM was predicted by LPMC RDM or MPMC RDM. In the MPMC model, Addressee RDM was also predicted by LMTG RDM, while the mean estimate of the model coefficient of MPMC RDM was numerically higher than that of LMTG RDM (0.3 vs. 0.11). In contrast, neither Speaker RDM nor Context RDM was predicted by any of the brain RDMs (*Table S5* and *Figure S6*). For “*Request vs. Reply-2*”, Speaker RDM was predicted by LPMC RDM. In contrast, neither Addressee RDM nor Context RDM were predicted by any of the brain RDMs (*Table S*5 and *Figure S6*).

Thus, in an extension of the MVPC results, the RSA results suggested that, relative to the perisylvian regions, the premotor cortex more robustly represents *Promise*-related information that is predicted by the addressee’s attitudes, and *Request*-related information that is predicted by the speaker’s attitudes.

### Results of the lesion study

To examine if the premotor cortex plays a causal role in comprehending linguistic communications, we conducted a lesion study in which we tested whether patients with lesions in the premotor cortex (see Figure 4a for their lesion locations) would have impairments in the comprehension task relative to healthy controls and lesion controls who had lesions not located in the premotor cortex (see Figure 4b and *Methods* for their lesion locations). Participants from the three groups were instructed to judge who would perform the action described in the critical sentence (i.e., the performer judgement), to rate the interlocutors’ attitudes, and to respond to comprehension questions in catch trials. The accuracies of the performer judgement and the comprehension questions were comparable between the three groups. For the ratings of the interlocutors’ attitudes, to simplify the task and to accommodate the patients’ cognitive state, we instructed the participants to only rate the speaker’s will and the addressee’s will. These two ratings would be sufficient to reflect the goal-related information as the above factor analyses had shown that the three speaker/addressee-related features were attributed to the same accounting factor of the speaker/addressee’s attitudes.

**Figure 4.**
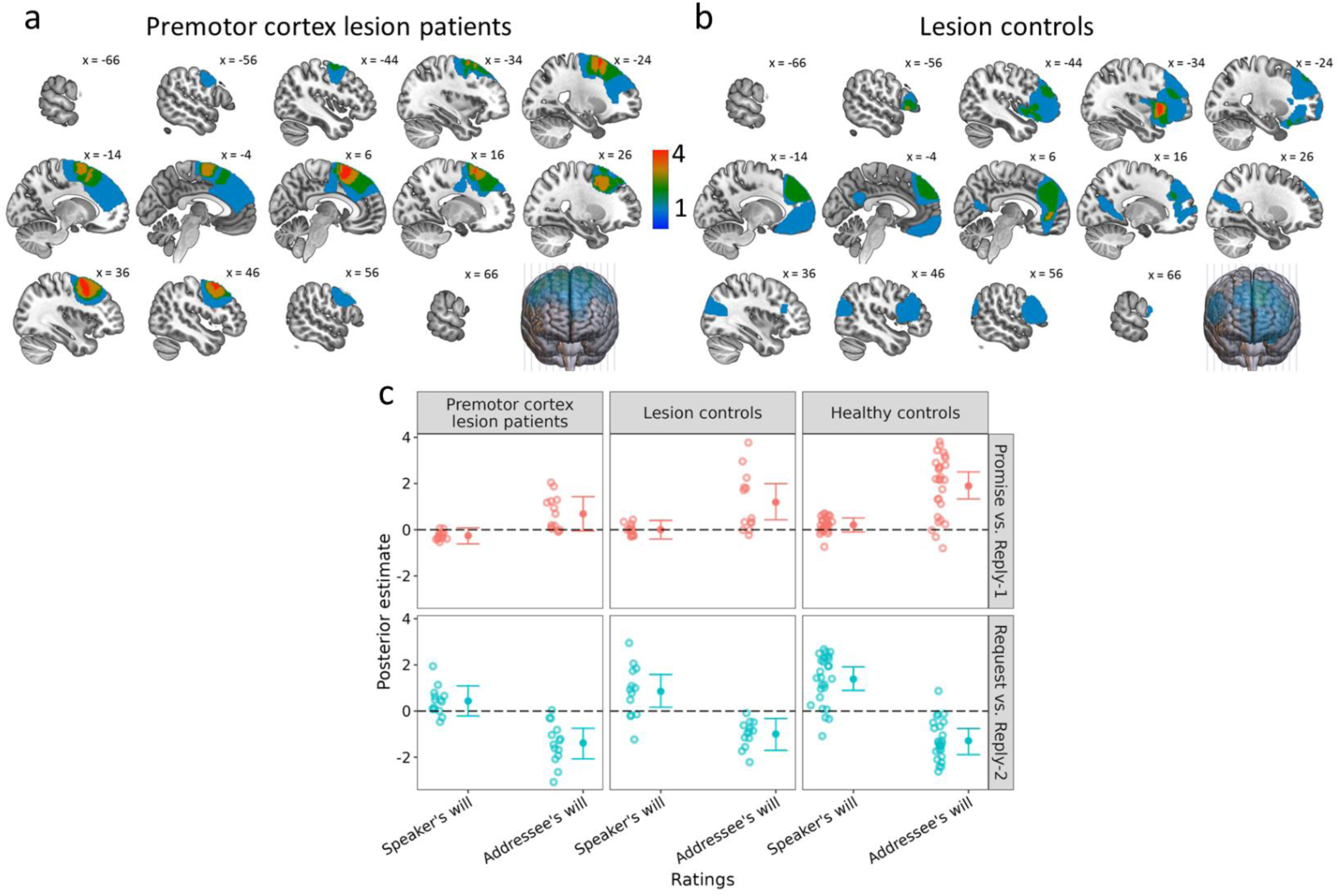
Results of the lesion study. Lesion reconstructions for **a,** premotor cortex lesion patients and **b,** lesion controls. The text in black indicates the x coordinates in the MNI system. The color bar indicates the number of patients. **c,** Results of the Bayesian hierarchical logistic model in the lesion study. The posterior estimates of the ratings (vertical axis) of the speaker’s will and the addressee’s will (horizontal axis) for the premotor lesion patients (left), lesion controls (middle), and healthy controls (right) are plotted. The upper panel represents the “*Promise* vs. *Reply-1*” model, the lower panel represents the “*Request* vs. *Reply-2*” model. The solid circles represent mean group-level posterior estimates. The error bars represent 95% *Crl*s. A 95% *CrI* excluding 0 indicates a significant group-level effect. The hollow circles on the left side of the group-level estimates represent the corresponding mean participant-level estimates for all the participants.

Bayesian hierarchical logistic models were fitted to estimate parameters at both the group-level and the participant-level. The group level estimates examined the extent to which communicative goals could be predicted by the ratings of the speaker’s will and the addressee’s will within each group (*Table S4*), and the participant-level estimates were fitted to test the difference in the predictability of the ratings between the groups. For “*Promise* vs. *Reply-1*” model (Figure 4c upper panel), the group-level results showed that communicative goals were significantly predicted by the addressee’s will rating for both lesion controls and healthy controls (*b* = 1.19, 95% *CrI*: [0.43, 1.99]; *b* = 1.89, 95% *CrI*: [1.33, 2.5], respectively), but not for the premotor lesion patients (*b* = 0.68, 95% *CrI*: [-0.05, 1.43]). For all groups, no significant predictability of the speaker’s will rating was observed. For the differences between the groups in the predictability of the addressee’s will rating, pair-wise comparisons^i^ on the participant-level estimates were performed. The results showed that the predictability of the addressee’s will rating for the premotor lesion patients was significantly lower than that for healthy controls (0.69 vs. 1.89, *t*_(38.62)_ = -3.76, *Cohen’s d* = 1.13, *p* < 0.001), with Bayes factor *BF*_10_ = 13, suggesting strong evidence for this difference^ii^. In contrast, there were no reliable difference between the predictability of the addressee’s will ratings for lesion controls and for healthy controls (1.19 vs. 1.89, *t*_(24.38)_ = -1.61, *Cohen’s d* = 0.54, *p* = 0.12), with Bayes factor *BF*_10_ = 0.86. As the pattern of the participant-level estimates of the predictability of the addressee’s will ratings showed the lowest mean value for the premotor lesion patients and the highest mean value for healthy controls, further linear regression modelling was performed to test the linear trend of these estimates across the groups, with the premotor lesion patients, lesion controls, and healthy controls coded as 1, 2, and 3 respectively. The result showed a significant positive slope for the groups (*β* = 0.61, *t* = 3.21, *p* = 0.002, *BF*_10_ = 15.63), suggesting that the comprehensibility of communication goals for *Promise* (vs. *Reply-1*) can be increasingly predicted by the addressee’s will rating over the three groups.

For “*Request* vs. *Reply-2*” (Figure 4c lower panel), the results of the group-level estimates showed that communicative goals were significantly predicted by the speaker’s will rating for both lesion controls and healthy controls (*b* = 0.85, 95% *CrI*: [0.17, 1.59]; *b* = 1.38, 95% *CrI*: [0.89, 1.92], respectively), but not for the premotor lesion patients (*b* = 0.44, 95% *CrI*: [-0.21, 1.09]). For all groups, “*Request* vs. *Reply-2*” were predicted by the addressee’s will ratings. For the differences between the groups in the predictability of the speaker’s will rating, further comparisons on the participant-level estimates were performed. The results showed that the predictability of the speaker’s will rating for the premotor lesion patients was significantly lower than that for healthy controls (0.44 vs. 1.38, *t*_(38.4)_ = -3.64, *Cohen’s d* = 1.1, *p* < 0.001), with Bayes factor *BF*_10_ = 10.92, suggesting strong evidence for this difference. In contrast, there was no reliable difference between the predictability of the speaker’s will rating for lesion controls and that for healthy controls (0.85 vs. 1.38, *t*_(22.26)_ = -1.41, *Cohen’s d* = 0.48, *p* = 0.17), with Bayes factor *BF*_10_ = 0.73. As the pattern of the participant-level estimates of the predictability of the speaker’s will ratings showed the lowest mean value for the premotor lesion patients and the highest mean value for healthy controls, further linear regression modelling was performed to test the linear trend of these estimates across the groups. The result showed a significant positive slope for the groups (*β* = 0.48, *t* = 3.03, *p* = 0.004, *BF*_10_ = 10.29), suggesting that the comprehensibility of the communication goals for *Request* (vs. *Reply-2*) can be increasingly predicted by the speaker’s will rating over the three groups.

Taken together, these results suggested that patients with lesions in the premotor cortex were impaired in comprehending communicative goals as compared with healthy controls. In contrast, there was no reliable difference between patients with lesions in brain regions other than the premotor cortex and healthy controls.

## Discussion

Across two studies, our results consistently and convergently demonstrated the role of the premotor cortex in representing communicative goals. The MVPC results showed that both the premotor cortex and the perisylvian regions contain information on communicative goals, and the results of the RS encoding models showed that the goal-related activity in these areas could be predicted by the interlocutors’ attitudes. These findings suggest that understanding linguistic communicative goals involves both the premotor cortex and the perisylvian regions. Moreover, the results of combinatorial MVPCs showed that the premotor cortex is more sensitive than the perisylvian regions to communicative goals, and the results of the RS decoding model showed that the goal-related activity in the premotor cortex is more reliably related to the interlocutors’ attitudes than that in the perisylvian regions. These findings suggest that the premotor cortex is more pronounced in representing communicative goals than the perisylvian regions.

Furthermore, in the lesion study, the results of both the comparisons of the participant-level estimates between the groups and the linear trend analyses showed that, the predictability of the speaker/addressee’s will ratings has the lowest value for patients with premotor lesions, the medium value for lesion controls, and the highest value for healthy controls. These results suggest that the premotor cortex lesions have profoundly impaired the understanding of communicative goals, demonstrating a causal role of the premotor cortex in representing goals. Collectively, the current study supports the theoretical view (Austin, 1975; Searle, 1969; Wittgenstein, 1953) that linguistic communications are represented as goal-directed actions in the brain.

In an extension of previous fMRI studies on linguistic communications (Egorova *et al*., 2016; Feng *et al*., 2017; Feng *et al*., 2021; Shibata *et al*., 2011; van Ackeren *et al*., 2012), we used sentences that directly convey communicative goals that are independent of action-related semantics. Specifically, our results showed that the activation patterns in the premotor cortex could discriminate the goals even though the critical sentences (e.g., “I/You will analyze the survey data this week.” in Table 1) express the same action semantics (e.g., “analyzing the survey data”) across conditions. These goal-related representations that go beyond action semantics suggest that the premotor cortex represents the relation between the interlocutors and the information of the speaker’s sentence. This relation can be reflected by the interlocutors’ attitudes (e.g., the speaker/addressee’s will/cost-benefit/pleasure) towards the information of the sentence (Searle and Vanderveken, 1985). Specifically, the premotor cortex represents not simply what action would be conducted (e.g., the data analysis) but more importantly the extent to which the interlocutors would like to have the action accomplished. In agreement with this argument, the results of both the RSA and the lesion study demonstrated that the goal-related activity pattern in the premotor cortex was related to the specific interlocutor’s attitude that predicted the goal. Specifically, the premotor cortex was sensitive to the addressee’s attitude when discriminating *Promise* from *Reply-1* while to the speaker’s attitude when discriminating *Request* from *Reply-2*. In a general sense, the interlocutors and the speaker’s sentence can be deemed as the subjects and their objectified intentions of the action. Such a generalization echoes with a recent study showing that the training of motor actions improves the understanding of subject-object relations in sentences (Thibault *et al*., 2021), demonstrating the general role of the motor system in understanding relations between subjects and objects of actions.

To establish the relations between the interlocutors and the sentential information for linguistic communications, the premotor cortex can function in a way like the understanding of motor actions (Rizzolatti et al., 1996; Rizzolatti and Sinigaglia, 2016). It has been suggested that human premotor cortex contains neurons with “mirror” properties (Rizzolatti and Arbib, 1998; Rizzolatti et al., 1988) that enable similar activation patterns for action implementation and action observation (Avenanti et al., 2007; Molenberghs et al., 2012; Oosterhof et al., 2012; 2013; Rizzolatti et al., 2014). Specifically, the goal-related premotor activity patterns induced by observing others’ actions are similar to those induced by implementing actions (Oosterhof *et al*., 2012). The premotor cortex has been repeatedly shown to be involved in understanding motor action goals (e.g., reaching or grasping) (Cattaneo et al., 2010; Gallivan et al., 2011a; Gallivan *et al*., 2013; Gallivan et al., 2011b; Michael et al., 2014). During the understanding of the goal of an observed action, the premotor cortex supports a reactivation of the representation of relevant action program, a process termed as action/mental simulation (Gallese and Goldman, 1998; Jacob and Jeannerod, 2005; Jeannerod, 2001; Zwaan, 2016). Similarly, in understanding the communicative goal conveyed by language, a mental simulation occurs to enable the comprehender to build a model that integrates the goal with the interlocutors’ attitudes; this simulation clarifies the relation between the interlocutors and the speaker’s sentence (Zwaan, 2016).

In processing the speaker’s meaning in linguistic communication, the comprehender could, on the one hand, mentally simulate the speaker’s communicative action, and/or, on the other hand, infer the speaker’s meaning and goal using a “theory of mind” about the speaker’s (and possibly addressee’s) mental state as suggested by previous studies (Feng *et al*., 2021; van Ackeren *et al*., 2012). This use of “theory of mind” is generally supported by the increased activations in medial prefrontal cortex (MPFC) and temporo-parietal junction (TPJ) (Schurz et al., 2014). Although the current fMRI experiment did observe that these typical “theory of mind” regions represented communicative goals, the strength of the representations in these regions was weaker than that in the premotor cortex. Our findings thus support the notion that understanding communicative goals is a relatively primitive and spontaneous process of projecting one’s own experience on the other’s action that requires mental simulation rather than more effortful inference based on a “theory” about the other (Gallese and Goldman, 1998; Gordon, 1992). In many situations this primitive simulation process is sufficient to support the social functions of language (Gallese, 2008), including communicating with others (Garrod and Pickering, 2004).

The shared mechanisms of mental simulation for language comprehension and motor action understanding is likely the evolutionary consequence of communicating with each other in a symbolic or language-like manner. On the one hand, nonhuman primates have communicative actions involving symbolic vocalizations (e.g., monkeys’ alarm calls for predators, Arnold and Zuberbühler, 2006; Seyfarth et al., 1980; and chimpanzees’ referential grunt for food, Watson et al., 2015) and gestures (e.g., Bonobos and chimpanzees’ meaningful gestures for social outcomes, Graham et al., 2018). Tool-using that involves the premotor region (Gallivan *et al*., 2013) is a type of intellectual behavior shared by humans and nonhuman primates (Seed and Byrne, 2010). On the other hand, the connectivity between the left posterior temporal cortex and the left inferior frontal cortex in the perisylvian area is weaker in nonhuman primates’ brain than in the human brain (Balezeau et al., 2020; Friederici, 2009). Consequently, the premotor region, rather than the perisylvian regions, is a common neural substrate that supports both humans and nonhuman primates’ communicative actions with mental simulation, and is the footstone of the co-evolution of humans’ linguistic ability and tool-using/making skills (Arbib, 2011; Stout and Chaminade, 2012).

To conclude, while both the premotor cortex and the perisylvian language regions represent the information on communicative goals and the interlocutors’ attitudes, the premotor cortex represents more information than the perisylvian regions. Moreover, lesions in the premotor cortex result in impaired processing of linguistic communications. These findings demonstrated that the premotor cortex is necessary for comprehending communicative goals in language processing, supporting the theoretical view that linguistic communications are represented as goal-directed actions in the brain.

## Methods

### Study 1: fMRI study

#### Participants

Fifty-eight native Chinese speakers (30 females, mean age = 22 years, *SD* = 3, *range*: [18, 31]) with normal or corrected-to-normal vision participated the fMRI experiment. None of them reported a history of neurological or psychiatric disorders. Written informed consent was obtained from each participant prior to the experiment. Two participants were excluded from data analysis due to dropping out. This study was performed in accordance with the Declaration of Helsinki and was approved by the Committee for Protecting Human and Animal Subjects of the School of Psychological and Cognitive Sciences at Peking University.

#### Design and materials

Eighty quadruplets of Chinese scripts describing daily-life scenarios were created and selected (Table 1). Each quadruplet included four scripts. Each script started with a context of a communication between two interlocutors, a speaker and an addressee. Then a sentence “and they were communicating with each other” was included to introduce the pre-critical sentence “then A (i.e., the speaker) said to B (i.e., the addressee)” and the final critical sentence. The critical sentence was said by the speaker to convey a communicative goal.

Each of the four scripts in a quadruplet corresponded to one of four experimental conditions. A specific experimental condition was defined by the communicative goal conveyed by the critical sentence. Specifically, in the condition termed *Promise*, the context described a task that the addressee was supposed to but yet unable to complete for a particular reason and the speaker was capable of doing this task for the addressee. In the following conversation, the speaker said the critical sentence with the first-person subject “*I*” to express the intention to do the task. In the condition termed *Reply-1*, the context described the same task as the task in the *Promise* condition but specified that it was the speaker who was supposed to complete the task. In the following conversation, the speaker said the critical sentence with the first-person subject to state that the speaker would do the task. The *Reply-1* condition served as the control condition for the *Promise* condition in the way that the critical sentences in the two conditions were the same, but these two critical sentences conveyed different goals. In the condition termed *Request*, the context described the same task as above that the speaker was supposed to yet be unable to complete for a particular reason, and the addressee was capable of doing this task for the speaker. In the following conversation, the speaker said the critical sentence with the second-person subject “*You*” to express the intention to request the addressee to do the task. In the condition termed *Reply-2*, the context described the same task that the addressee was supposed to complete. In the following conversation, the speaker said the critical sentence with the second-person subject to describe that the addressee would do the task. The *Reply-2* condition served as the control condition for the *Request* condition in the way that the critical sentences in the two conditions were the same, but these two critical sentences conveyed different goals. The pre-critical sentence was consistent across the four conditions in a quadruplet because the identities of the speaker and the addressee were invariable. The critical sentences varied only in the subjects, resulting in identical critical sentences for *Promise* and *Reply-1* and for *Request* and *Reply-2* respectively.

These scripts were selected based on the evaluative results from an independent group of participants in a pilot rating task prior to the fMRI experiment (*Supplemental Information*).

The 80 quadruplets of scripts (320 scripts in total) were assigned into 4 experimental lists according to a Latin-square procedure. Each experimental list included 80 scripts, with 20 scripts for each condition. In a specific experimental list, 80 scripts came from the 80 different quadruplets such that there was no repetition of a specific critical sentence. For each participant, based on the Latin-square design, scripts from one of the experimental lists were used as the reading materials. The 80 scripts (trials) were divided into five scanning runs (16 scripts per run), each of which lasted approximately seven minutes. Trials of the four conditions were equally distributed in the five scanning runs. In each run, trials of different conditions were mixed and presented in a pseudo-randomized order with the restriction that no more than three scripts with the same communicative goal were presented consecutively.

Each participant was asked to silently read a list of scripts in the MR scanner. Stimuli were presented in black (RGB: 0, 0, 0) against a gray background (RGB: 180, 180, 180). In each trial (Figure 1a), a fixation cross was firstly presented at the center of the screen for a jitter duration of 1-5.5 s, followed by a cross presented at the upper left part of the screen where the first character of the context would located. This fixation was presented for 1 s to direct participants’ attention. The context was presented for 10 s and followed by a cross presented at the center for another jitter duration of 1-3.25 s. A cross was then presented for 1 s at the upper left part of the screen where the first character of the pre-critical sentence was located. After the offset of the cross, the pre-critical sentence was presented. After the pre-critical sentence had been presented for 1 s, a cross was presented below the pre-critical sentence, where the first character of the critical sentence would be located. This cross lasted for 0.5 s together with the pre-critical sentence. After the offset of the cross, the critical sentence was presented within double quotes for 4 s, together with the pre-critical sentence. To engage the participants into the script reading, a comprehension question was added to each of the 24 catch trials (30% of the all trials). At the end of these catch trials, a triangle was presented at the center for a jitter duration of 1-6.6 s, followed by a comprehension question. Participants were instructed to make “yes” or “no” response by pressing the button on the response box in their left or right hand. Half of the participants were instructed to press the left button for “yes” and the right button for “no”, and the other half made their responses with a reversed button-hand assignment. Half of these trials required a “yes” as correct response and the other half required a “no” as correct response.

Prior to the scanning, participants performed ten practice trials with scripts not in the experimental lists. After the scanning, participants were asked to fulfill a post-scanning rating task on the same scripts they read in the scanner. They were asked to rate with 7-point scales on the following ten features: speaker’s will, speaker’s cost-benefit, speaker’s pleasure, addressee’s will, addressee’s cost-benefit, addressee’s pleasure, social distance between the interlocutors, relative power between the interlocutors, the performer’s capability, and the mitigation of the critical sentence. These ten features were the same as those rated in the pilot evaluation (see *Supplemental Information*).

#### Statistical analysis of post-scanning ratings

##### Bayesian logistic mixed models

To assess the extent to which communicative goals could be predicted by the ten features, the post-scanning ratings were fitted with Bayesian logistic mixed models using the *brms* package (Bürkner, 2017) in R environment. The two pair-wise predictions, “*Promise* vs. *Reply-1*” and “*Request* vs. *Reply-2*”, were assessed respectively with an independent model. In each model, the response variable was the communicative goal, the predictors were the ratings of the ten features. Full models were fitted to reduce type-I error rate (Barr et al., 2013). The priors for all fixed slopes and fixed intercept were *Normal*(0,100), while the priors for standard deviations were *Cauchy*(0,5). Within the variance-covariance matrices of the by-participant and by-item random effects, priors were defined for the correlation matrices using a LKJ prior with parameter *η* 1.0(Lewandowski et al., 2009). The joint posterior distribution was sampled by four Monte-Carlo Markov Chains (MCMCs) at 20,000 iterations for each model, with the first half of the samples discarded as warm-up samples. Convergence was checked using 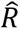 convergence diagnosis (Gelman and Rubin, 1992). Mean estimates (*b*) and 95% credible intervals (*CrI*s) of posterior distributions were used to evaluate the fitted coefficients. All posterior estimates reported have 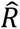-values lower than 1.01. The predictability of the contextual features was indexed by the corresponding posterior estimate, and the estimates were considered as statistically significant when the 95% *CrI* excluded 0.

##### Factor analyses

To evaluate the reliability of the three-factor model estimated by the EFA on the post-scanning ratings (*Supplemental Information*), a confirmatory factor analysis (CFA) with the factor structure obtained by the EFA was conducted using the *lavaan* package(Oberski, 2014) in R. The CFA model was evaluated by *CFI*, *TLI*, and *RMSEA*.

#### MRI data acquisition and preprocessing

A GE-MR750 3T MR scanner was used to collect T1-weighted structural images with 1 × 1 × 1 mm^3^ voxel size and functional images. In each run of fMRI, an echo-planar imaging (EPI) sequence with an interleaved (bottom-up) acquisition order, 2,000 ms repetition time, 30 ms echo time, and 90° flip angle to obtain 225 3D volumes of the whole brain. Each volume consisted of 33 axial slices covering the whole brain. Slice thickness was 3.5 mm and inter-slice gap was 0.7 mm, with a 224 mm field of view (FOV), 64 × 64 matrix, and 3.5 × 3.5 × 4.2 mm^3^ voxel size. Head motion was minimized using cushions around the head and a forehead strap.

The preprocessing of fMRI data was implemented using FEAT (fMRI Expert Analysis Tool) in FSL (FMRIB Software Library v5.0.11) (Jenkinson et al., 2012). To ensure steady state magnetization, the first five volumes were discarded. Preprocessing consisted of brain extraction using BET based on the structural image (Smith, 2002), motion correction using MCFLIRT (Jenkinson et al., 2002), slice-timing with Fourier-space time-series phase-shifting, spatial smoothing with a Gaussian kernel of FWHM 5mm, high-pass temporal filtering with a cutoff of 100 s, and grand-mean intensity normalization of the entire 4D dataset by a single multiplicative factor. These preprocessing procedures were applied on fMRI data for both univariate and multivariate analyses, except that spatial smoothing was skipped for the multivariate analyses. To co-register the structural image and the functional images, a linear transformation with 12 degrees of freedom (*df*) allowing translation and rotation was applied by FLIRT (Jenkinson *et al*., 2002; Jenkinson and Smith, 2001). The 12 *df* linear transformation from the structural image to the MNI system was further refined using FNIRT registration with nonlinear algorithm for 54 participants’ images, whereas the registration with linear algorithm was used for the other two participants’ images for better alignment.

#### ROIs definitions

The bilateral premotor cortex, including medial premotor cortex (MPMC) and lateral premotor cortex (LPMC), were defined based on the probability maps of the Jülich Histological atlas (Geyer, 2004). In the premotor cortex (Figure 2a), voxels with an absolute value of x coordinate in the MNI system lower than 16 were assigned to MPMC, otherwise, they were assigned to LPMC (Mayka *et al*., 2006).

Four ROIs in the left lateralized perisylvian language region, including BA44 division of the Broca’s area (left BA44), BA45 division of the Broca’s area (left BA45) (Amunts et al., 1999), left middle temporal gyrus (LMTG), and left superior temporal gyrus (LSTG) (Desikan et al., 2006) were defined based on the probability maps of the Jülich Histological atlas and the Harvard-Oxford Cortical atlas (Figure 2a).

For all ROIs, only voxels with a probability greater than 20% were reserved. Each voxel was assigned to a ROI according to its maximum probability among the above mentioned five ROI probability maps.

#### General linear model (GLM)

For each participant, four regressors were respectively defined for the four experimental conditions, *Promise*, *Reply-1*, *Request*, and *Reply-2*, by the onsets of the critical sentence with a duration of 4 s convolved by canonical hemodynamic response function (HRF). The temporal derivative of each of the four regressors was included as a regressor of no-interest. Another three regressors respectively corresponding to the context, the pre-critical sentence, and the comprehension question were also included. The six parameters of head movements were added in to reduce the influence of head motion on signal changes.

#### ROI-based multivariate pattern classification (MVPC)

To detect differences in activity patterns representing the different communicative goals, MVPC was conducted for each ROI using the *PyMVPA* toolbox (Hanke et al., 2009). In each ROI, the voxel-level parameter estimates of the critical sentences were extracted, detrended along time series, and transformed to *Z*-scores across runs. For each ROI, cross-validated classifications of communicative goals were performed using a linear support vector machine (SVM) as a classifier. Four pair-wise classifications were performed: (1) *Promise* vs. *Reply-1*; (2) *Request* vs. *Reply-2*; (3) *Promise* vs. *Request*; (4) *Reply-1* vs. *Reply-2*. For each pair-wise classification, a participant-based cross-validation with 50 repetitions were conducted. Each repetition consisted of a training set of data from 45 (approximately 80% of all data) randomly selected participants and a test set of data from the remaining 11 participants (approximately 20% of all data). For each repetition, a cross-validated accuracy was computed as a percentage of correct classifications of the test set to evaluate the performance of a classifier, and the mean accuracy averaged over the 50 repetitions was calculated.

Statistical significance of the classification accuracy was tested using permutation-based classifications with 2,000 repetitions for each pair-wise classification (Stelzer et al., 2013). In each repetition, the participant-based cross-validation procedure described above was performed on the data with permuted communicative goals, generating 2,000 null cross-validated accuracies derived for each pair-wise classification. Probabilities (*p*-values) of the observed accuracies against the distribution of the permutation-based null accuracies were computed. Statistical significance was determined by a Bonferroni-corrected significance threshold of *p* < 0.002 (24 comparisons were conducted in total).

##### Combinatorial MVPC

To examine whether the lateral/medial premotor cortex represented more information on communicative goals relative to the ROIs in the perisylvian region, combinatorial MVPCs (Carter *et al*., 2012; Clithero *et al*., 2009) were conducted. The current analyses focus on two pair-wise classifications, “*Promise* vs. *Reply-1*” and “*Request* vs. *Reply-2*”.

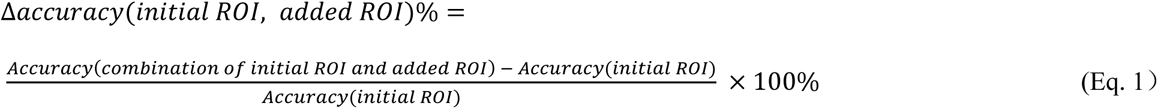

The computation is performed by Eq. 1. Combinatorial accuracy, *Accuracy(combination of initial ROI and added ROI),* was the cross-validated accuracy based on voxels collapsed over an initial ROI and an added ROI. *Δaccuracy(initial ROI, added ROI)* was obtained by subtracting the cross-validated accuracy based on the voxels in an initial ROI, i.e., *Accuracy(initial ROI),* from the combinatorial accuracy, and dividing this difference by the *Accuracy(initial ROI)*.

This *Δaccuracy(initial ROI, added ROI)* is an index to quantify the extent to which the added ROI improved classification performance based on the initial ROI. These allow us to examine improvements in classification accuracy contributed by a premotor ROI for each of the perisylvian ROIs and vice versa.

Two steps of permutation-based significance testing were conducted on *Δaccuracy(initial ROI, added ROI)*:

(1) To test whether the premotor ROIs and the perisylvian ROIs improved classification performance to each other, we conducted 32 (2 premotor ROIs × 4 perisylvian ROIs × 2 pair-wise classifications × 2 alternatives of initial-added ROIs pair) permutation tests with 2,000 repetitions. For each test, 50 cross-validated accuracies of the initial ROI and 50 cross-validated combinatorial accuracies served as observations. These two types of cross-validated accuracies were permuted and used to compute a null *Δaccuracy(initial ROI, added ROI)* every repetition, generating a set of null *Δaccuracy(initial ROI, added ROI)*, and *p*-value for the observed accuracy against the null distribution was computed. Statistical significance was determined by Bonferroni-corrected significance threshold of *p* < 0.0016.
(2) To test whether the premotor ROIs represented more information on communicative goals relative to the perisylvian ROIs, we conducted 16 (2 premotor ROIs × 4 perisylvian ROIs × 2 pair-wise classifications) pair-wise permutation tests with 2,000 repetitions to compare *Δaccuracy(an ROI in perisylvian region, lateral/medial premotor cortex)* with *Δaccuracy(lateral/medial premotor cortex, an ROI in perisylvian region)*. For each test, the two types of *Δaccuracies* were permuted every repetition to generate a set of null differences between *Δaccuracy(an ROI in perisylvian region, lateral/medial premotor cortex)* and *Δaccuracy(lateral/medial premotor cortex, an ROI in perisylvian region)*, and *p*-value of the observed difference against the null distribution was computed. Statistical significance was determined by a Bonferroni-corrected significance threshold of *p* < 0.003.

#### Representational Similarity Analyses (RSAs)

RSAs (Kriegeskorte *et al*., 2008) were conducted to further examine if the six ROIs represent the information on the speaker’s attitudes and the addressee’s attitudes, and if the premotor ROIs represented more information on the speaker/addressee’s attitudes relative to the perisylvian ROIs. These analyses were implemented for two pair-wise predictions, “*Promise* vs. *Reply-1*” and “*Request* vs. *Reply-2*”, respectively.

For each participant, the GLM was refitted with the same definitions of regressors as described above, except that the critical sentence of every trial was defined as a single regressor. Hence, each GLM had 80 regressors of the critical sentences. For each ROI of each participant, to avoid over-fitting, feature selection was conducted on the voxels of each ROI(Hanke *et al*., 2009), and voxels with the 50% highest F values were included in the RSA. For the selected voxels, parameter estimates of the critical sentences from the GLM were extracted and transformed to Z-scores. Pattern similarity matrix of each ROI was built up by calculating the Pearson correlation of the voxel-wise Z-scores between each two trials. Then the representational dissimilarity matrix (RDM) was obtained by 1 – similarity matrix, resulting in a 40 (2 conditions * 20 trials per condition) * 40 brain RDM for each ROI of each participant. For each participant, based on the post-scanning ratings and the three-factor model obtained by the factor analyses, independent behavioral RDMs were built up to represent the variance of the speaker’s attitudes (Speaker RDM), the variance of the addressee’s attitudes (Addressee RDM), and the variance of the contextual information (Context RDM), respectively. For each behavioral factor, the ratings of the three dimensions for each trial were represented in a three-dimensional space, and the Euclidian distance between each two rating points was calculated as the value in each cell of the behavioral RDM. Each of the three behavioral RDM also had a 40 (2 conditions * 20 trials per condition) * 40 structure.

For each ROI and each of the two pair-wise predictions, a representational similarity (RS) encoding model was used to assess the extent to which the brain RDM can be predicted by the behavioral RDMs. Specifically, a Bayesian linear mixed model was conducted with Eq. 2, where the brain RDM was included as the response variable and the three behavioral RDMs as the predictors. In each model, the response variable *Brain_RDM_i_* indicates the brain RDM of the *i*th participant (*i* ∊ [1, 56]). The predictor *Behavioral_RDM_k_* indicates one of the behavioral RDMs (*k* ∊ [1, 3]). Fixed effects consisted of the fixed slopes *b_k_* and the fixed intercept *b_o_*. Random effects consisted of by-participant random slope *v_ik_* for the *k*th behavioral RDM and random intercept *v_i_*_0_ for the *i*th participant. The model also included the *i*th participant’s residual *e_i_*. Only the lower triangular RDMs were used in these analyses. The model-fitting method was identical to the analysis of the post-scanning ratings. All posterior estimates reported have 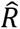 lower than 1.01.

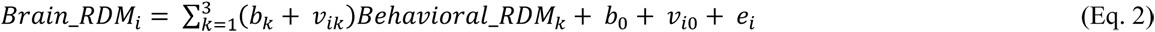

For each of the two pair-wise predictions and each of the behavioral RDMs, a RS decoding model was used to assess the extent to which the behavioral RDM can be predicted by the brain RDMs. Specifically, a Bayesian linear mixed model was conducted with Eq. 3, where the behavioral RDM was included as the response variable and the RDMs of five ROIs as the predictors.

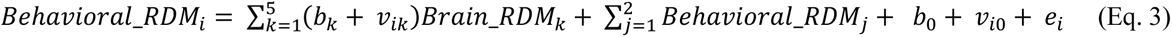

In each model, the response variable *Behavioral_RDM_i_* indicates the behavioral RDM of the *i*th participant (*i* ∊ [1, 56]). The predictors *Brain_RDM_k_* indicate one of the five brain RDMs (*k* ∊ [1, 5]), which were RDMs of the four perisylvian ROIs and an RDM of one of the premotor ROIs (the RDMs of LPMC and MPMC were included in distinct models respectively to prevent co-linearity). To regress out variances of the two behavioral RDMs other than the *Behavioral_RDM_i_*, they were included in the model as covariates *Behavioral_RDM_j_* (*j* = 1, 2). The meaning of the remaining parameters in Eq. 3 were the same as the RS encoding model. The model-fitting method was identical to the analyses of the post-scanning ratings. All posterior estimates reported have 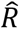 lower than 1.01.

As described above, while the RS encoding model estimated the coefficients of the different behavioral RDMs in predicting the brain RDM of a specific ROI, the RS decoding model estimated the coefficients of the brain RDMs of different ROIs in predicting a specific behavioral RDM. An effect was tested with Bonferroni corrected *CrI*s, 99.86% *CrI* for the encoding models where 24 (6 brain RDMs × 3 behavioral RDMs × 2 pair-wise predictions) effect estimates were tested, and 99.92% *CrI* for the decoding models where 60 (3 behavioral RDMs × 5 brain RDMs × 2 models with different premotor ROIs × 2 pair-wise predictions) effect estimates were tested. An effect was determined as significant when Bonferroni corrected *CrI* excluded 0.

### Study 2: lesion study

#### Participants

Twenty-nine adult patients with unilateral lesions recruited from the Patient’s Registry of Beijing Tiantan Hospital (Beijing, China) participated in the experiment. Their lesions resulted from the surgical removal of low-grade gliomas. Depending on the lesion locations, the patients were assigned to either the premotor cortex lesion group or lesion control group. Thirty healthy adults without a known history of psychiatric or neurological disorder were recruited from the local community as a healthy control group. All participants are native Chinese speakers with normal or corrected-to-normal vision and right-handed, with handedness assessed by the Chinese Handedness Questionnaire (Li, 1983). Two lesion control participants and three healthy control participants were excluded from the data analyses due to poor task performances (see below). This study was performed in accordance with the Declaration of Helsinki and was approved by the Committee for Protecting Human and Animal Subjects of the School of the Psychological and Cognitive Sciences at Peking University and the Institutional Review Board of the Beijing Tiantan Hospital at Capital Medical University.

The demographic variables of the participants are shown in *Table S*6. The comparisons of the demographic variables were conducted by independent-sample *t*-tests. The lesion sizes and the chronicities were comparable between the premotor lesion patients and lesion controls (all *p*-values > 0.2). Participants’ mental states were assessed with the Mini-Mental State Examination (MMSE) (Hamilton et al., 1976) and the Beck depression inventory (BDI) (Knight, 1984). The MMSE scores and the BDI scores were comparable between all groups (all *p*-values > 0.1). Years of education were comparable between the premotor lesion patients and lesion controls (*p* = 0.734). The years of education of healthy controls were slightly higher than the premotor lesion patients and lesion controls, but did not reach the significance level (*p* = 0.057; *p* = 0.065, respectively). While the participants’ ages were comparable between the premotor lesion patients and lesion controls and between the premotor lesion patients and healthy controls (*p*-values > 0.1), lesion controls were older than healthy controls in average (*t*_(18.1)_ = 2.58, *p* = 0.019).

#### Lesion reconstruction and group assignment for patients

Two neurosurgeons (the third author and the forth author) identified the lesions of each patient and created the lesion masks based on the structural images. The lesion masks were transformed into the MNI system by linear registration using FSL. For each patient, there were four steps of the registration. First, based on T1-weighted structural image, white matter mask was extracted using the *fast* function (Zhang et al., 2001). Second, based on the T1-weighted image, the white matter mask, and the lesion mask, the lesion area in the T1-weighted image was filled using the *lesion_filling* function (Battaglini et al., 2012). Third, the filled T1-weighted image was transformed into the MNI system by linear registration. This registration generated a transformed matrix of the spatial relation between the filled T1-weighted image and the MNI system. Finally, based on the transformed matrix, the lesion mask was transformed into the MNI system. To ensure the results of the registrations were consistent with the patients’ clinical situation, the neurosurgeons further checked and modified the transformed lesion masks. We computed the overlapped volume between each transformed lesion mask and the premotor cortex probability maps of the Jülich Histological atlas. Fourteen patients with overlapped volumes larger than 2 ml were assigned to the premotor lesion group (Figure 4a), other 15 patients were assigned to the lesion control group (Figure 4b). In the lesion control group, four patients had lesions in the left prefrontal cortex, four in the right prefrontal cortex, four in the left insula, two in the right temporal cortex, and one in the right parietal cortex.

#### Design and procedure

Eighty-four quadruplets of scripts were created. The structure of the scripts and the experimental design were the same as the fMRI study with the exception that the contents of the scripts were easier to understand to accommodate the patients’ cognitive states. To evaluate the reliability of the scripts, two pilot studies with independent groups of participants were conducted beforehand (*Supplemental Information*). We firstly evaluated the scripts and replicated the pattern of results in the fMRI post-scanning ratings, and secondly assessed the appropriateness of the experimental procedure (see below). The results indicated that healthy adults were able to understand the scripts and to complete the task following instructions (*Supplemental Information*).

The scripts were assigned into four experimental lists according to a Latin-square procedure. Each list included 84 scripts (trials), and each list was further divided into four sections corresponding to four experimental blocks. Each participant was presented a list of scripts with a pseudo-randomized order. No more than three scripts with the same communicative goal were presented consecutively.

The experiment began with ten practice trials, followed by the four blocks of the main experiment. Each block began with a warm-up trial. The scripts used for the practice trials and the warm-up trials were not in the experimental lists. Each trial of the main experiment had the same sequence of the events of the script presentations as the fMRI experiment, except that the duration of each event was longer to accommodate the patients’ cognitive state (*Figure S*1). After the presentation of the critical sentence, participants were instructed to respond to three or four questions. First, they were instructed to judge who would perform the action described in the critical sentence (i.e., the performer judgement) within 10 s. The names of the speaker and the addressee were randomly presented on the left bottom and the right bottom of the screen respectively, participants had to choose either of the names by pressing the button on the corresponding side. Second, they were instructed to rate the speaker’s will and the addressee’s will on a 7-point scale, each of which had to be completed within 20 s. To accommodate the patients’ cognitive state, the whole script was presented at the top of the screen to allow them to reread the script. The two ratings were presented in random order. To engage participants into the reading, 24 catch trials (29% of all trials) with comprehension questions regarding the scripts were included. In each catch trial, a triangle was presented at the center after the ratings for a jitter duration of 0.5-1.5 s followed by a Yes/No comprehension question, and participants were asked to answer the comprehension question by pressing the corresponding button.

#### Data analyses

Two lesion control participants, who had lesions in the left insula and left temporal cortex respectively, and three healthy control participants were excluded from the data analyses because their accuracies for either the performer judgements or the comprehension questions were below two standard deviations from the mean.

##### Bayesian hierarchical logistic models

To assess the extent to which communicative goals could be predicted by the speaker’s will and the addressee’s will, we fitted Bayesian hierarchical logistic models for “*Promise* vs. *Reply-1*” and “*Request* vs. *Reply-2*” respectively using *Stan* (Carpenter et al., 2017) in R. To exclude trials with scripts that were apparently not comprehended by the participants, the model-fitting only used the trials with correct performer judgements, leaving 91% of the data. The communicative goal was included as the response variable, the ratings of the speaker’s will and of the addressee’s will were included as the predictors, and the age of the participants was included as a covariate to regress out the variance of the age, as shown in Eq. 4.

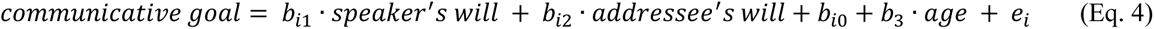

In each hierarchical model, the slopes of the speaker’s will rating and the addressee’s will rating and the intercept were estimated at the group-level and the participant-level, respectively. The group-level analysis independently estimated the parameters for each group and tested the effect estimates of the ratings within each of the three independent groups. The participant-level analysis independently estimated the parameters for each participant and tested the difference in the estimates between the groups. Each group-level parameter had a normal prior distribution, which had a mean with a prior of *Normal(0,100)* and a standard deviation with a prior of *Cauchy(0,5)*. Each of the participant-level parameters also had a normal prior distribution, which had a mean equaling to the mean of the corresponding group-level distribution and a standard deviation with a prior of *Cauchy(0,5)*. These parameters were shown in Eq. 4, for the *i*th participant, with the slopes of the two ratings, *b_i_*_1_ and *b_i_*_2_, and the intercept, *b_i_*_0_. The model also included *b*_3_ as the slope of the participants’ age and *e_i_* as the *i*th participant’s residual. Each of these parameters had a normal prior distribution, which had a mean with a prior of *Normal(0,100)* and a standard deviation with a prior of *Cauchy(0,5)*. The model-fitting method was the same as the analysis of the ratings in the fMRI study. All posterior estimates reported have 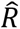 lower than 1.01.

Furthermore, we conducted Frequentist *t*-tests and Bayesian *t*-tests to compare the participant-level posterior estimates of the ratings between the groups. T-statistics, *Cohen’s d*-values, *p*-values, and Bayes factors in supporting H_1_ against H_0_, *BF*_10_, are reported in *Results*. To test the linear trend of the participant-level estimates across groups, we fitted linear regression models that included these estimates as response variables and the group as the predictor, with the premotor lesion patients coded as 1, lesion controls coded as 2, and healthy controls coded as 3. The slopes for the groups, *b*, the corresponding *t*-statistics, *p*-values, and the Bayes factors *BF*_10_ against the intercept-only models are reported in *Results*.

## Acknowledgement

This work was sponsored by the China Postdoctoral Science Foundation (2021M702211, awarded to W.C.), the Shanghai Sailing Program (20YF1422100, awarded to L.W.), the National Science Foundation of China (32000779, awarded to L.W.), the National Natural Science Foundation of China (81729001, awarded to Z.G.), and the National Science Foundation of China (31630034, 71942001, awarded to X.Z.). We thank Drs. Yingying Tan and Xiaoming Jiang for their comments on an earlier version of the manuscript. The data analyses were conducted on the High-performance Computing Platform of Peking University.

## Author Contributions

W.C. and X.Z designed the experiments. W.C., R.Y., X.W., and Z.G. conducted the experiments. W.C. and L.W. analyzed the data. W.C., L.W., and X.Z. wrote the paper. R.Y., X.W., and Z.G. revised the paper.

## Footnotes

i Each comparison of the participant-level posterior estimates between two groups tested the null hypothesis (H_0_), “there was no difference in the estimates between the groups”, against the alternative hypothesis (H_1_), “there was a difference in the estimates between the groups”. See *Methods* for details.

ii According to Andraszewicz and colleagues’ suggestion (2015), a *BF*_10_ between 10 and 30 indicates strong evidence for H_1_, and a *BF*_10_ between 0.33 and 1 indicates anecdotal evidence for H_0_.

## References

Amunts, K., Schleicher, A., Bürgel, U., Mohlberg, H., Uylings, H.B.M., and Zilles, K. (1999). Broca’s region revisited: Cytoarchitecture and intersubject variability. Journal of Comparative Neurology 412, 319–341. 10.1002/(sici)1096-9861(19990920)412:2<319::Aid-cne10>3.0.Co;2-7.

Andraszewicz, S., Scheibehenne, B., Rieskamp, J., Grasman, R., Verhagen, J., and Wagenmakers, E.-J. (2015). An introduction to Bayesian hypothesis testing for management research. Journal of Management 41, 521–543. 10.1177/0149206314560412.

Arbib, M.A. (2011). From mirror neurons to complex imitation in the evolution of language and tool use. Annual Review of Anthropology 40, 257–273. 10.1146/annurev-anthro-081309-145722.

Arbib, M.A. (2016). Towards a Computational Comparative Neuroprimatology: Framing the language-ready brain. Physics of Life Reviews 16, 1–54. https://doi.org/10.1016/j.plrev.2015.09.003.

Arnold, K., and Zuberbühler, K. (2006). Semantic combinations in primate calls. Nature 441, 303–303. 10.1038/441303a.

Austin, J.L. (1975). How to do things with words (Harvard University Press).

Avenanti, A., Bolognini, N., Maravita, A., and Aglioti, S.M. (2007). Somatic and Motor Components of Action Simulation. Current Biology 17, 2129–2135. 10.1016/j.cub.2007.11.045.

Aziz-Zadeh, L., Koski, L., Zaidel, E., Mazziotta, J., and Iacoboni, M. (2006). Lateralization of the human mirror neuron system. The Journal of Neuroscience 26, 2964–2970. 10.1523/jneurosci.2921-05.2006.

Balezeau, F., Wilson, B., Gallardo, G., Dick, F., Hopkins, W., Anwander, A., Friederici, A.D., Griffiths, T.D., and Petkov, C.I. (2020). Primate auditory prototype in the evolution of the arcuate fasciculus. Nature Neuroscience 23, 611–614. 10.1038/s41593-020-0623-9.

Barr, D.J., Levy, R., Scheepers, C., and Tily, H.J. (2013). Random effects structure for confirmatory hypothesis testing: Keep it maximal. Journal of Memory and Language 68, 255–278. https://doi.org/10.1016/j.jml.2012.11.001.

Battaglini, M., Jenkinson, M., and De Stefano, N. (2012). Evaluating and reducing the impact of white matter lesions on brain volume measurements. Human Brain Mapping 33, 2062–2071. 10.1002/hbm.21344.

Bianchi, S., Reyes, L.D., Hopkins, W.D., Taglialatela, J.P., and Sherwood, C.C. (2016). Neocortical grey matter distribution underlying voluntary, flexible vocalizations in chimpanzees. Scientific Reports 6, 34733. 10.1038/srep34733.

Brennan, S.E., Galati, A., and Kuhlen, A.K. (2010). Chapter 8 - Two minds, one dialog: Coordinating speaking and understanding. In Psychology of Learning and Motivation, B.H. Ross, ed. (Academic Press), pp. 301–344. 10.1016/S0079-7421(10)53008-1.

Bürkner, P.-C. (2017). brms: An R Package for Bayesian Multilevel Models Using Stan. Journal of Statistical Software 80, 1–28. 10.18637/jss.v080.i01.

Carpenter, B., Gelman, A., Hoffman, M.D., Lee, D., Goodrich, B., Betancourt, M., Brubaker, M., Guo, J., Li, P., and Riddell, A. (2017). Stan: A probabilistic programming language. Journal of Statistical Software 76, 1–32. 10.18637/jss.v076.i01.

Carter, R.M., Bowling, D.L., Reeck, C., and Huettel, S.A. (2012). A distinct role of the temporal-parietal junction in predicting socially guided decisions. Science 337, 109–111. 10.1126/science.1219681.

Cattaneo, L., Barchiesi, G., Tabarelli, D., Arfeller, C., Sato, M., and Glenberg, A.M. (2010). One’s motor performance predictably modulates the understanding of others’ actions through adaptation of premotor visuo-motor neurons. Social Cognitive and Affective Neuroscience 6, 301–310. 10.1093/scan/nsq099.

Clithero, J.A., Carter, R.M., and Huettel, S.A. (2009). Local pattern classification differentiates processes of economic valuation. NeuroImage 45, 1329–1338. https://doi.org/10.1016/j.neuroimage.2008.12.074.

Courson, M., Macoir, J., and Tremblay, P. (2017). Role of medial premotor areas in action language processing in relation to motor skills. Cortex 95, 77–91. 10.1016/j.cortex.2017.08.002.

Desikan, R.S., Ségonne, F., Fischl, B., Quinn, B.T., Dickerson, B.C., Blacker, D., Buckner, R.L., Dale, A.M., Maguire, R.P., Hyman, B.T., et al. (2006). An automated labeling system for subdividing the human cerebral cortex on MRI scans into gyral based regions of interest. NeuroImage 31, 968–980. https://doi.org/10.1016/j.neuroimage.2006.01.021.

Dreyer, F.R., and Pulvermüller, F. (2018). Abstract semantics in the motor system? – An event-related fMRI study on passive reading of semantic word categories carrying abstract emotional and mental meaning. Cortex 100, 52–70. 10.1016/j.cortex.2017.10.021.

Egorova, N., Shtyrov, Y., and Pulvermüller, F. (2016). Brain basis of communicative actions in language. NeuroImage 125, 857–867. 10.1016/j.neuroimage.2015.10.055.

Feng, W., Wu, Y., Jan, C., Yu, H., Jiang, X., and Zhou, X. (2017). Effects of contextual relevance on pragmatic inference during conversation: An fMRI study. Brain and Language 171, 52–61. 10.1016/j.bandl.2017.04.005.

Feng, W., Yu, H., and Zhou, X. (2021). Understanding particularized and generalized conversational implicatures: Is theory-of-mind necessary? Brain and Language 212, 104878. https://doi.org/10.1016/j.bandl.2020.104878.

Friederici, A.D. (2009). Pathways to language: fiber tracts in the human brain. Trends in Cognitive Sciences 13, 175–181. https://doi.org/10.1016/j.tics.2009.01.001.

Friederici, A.D. (2011). The brain basis of language processing: From structure to function. Physiological Reviews 91, 1357–1392. 10.1152/physrev.00006.2011.

Friederici, A.D., Chomsky, N., Berwick, R.C., Moro, A., and Bolhuis, J.J. (2017). Language, mind and brain. Nature Human Behaviour 1, 713–722.

Gallese, V. (2008). Mirror neurons and the social nature of language: The neural exploitation hypothesis. Social Neuroscience 3, 317–333. 10.1080/17470910701563608.

Gallese, V., and Goldman, A. (1998). Mirror neurons and the simulation theory of mind-reading. Trends in Cognitive Sciences 2, 493–501. 10.1016/S1364-6613(98)01262-5.

Gallese, V., and Lakoff, G. (2005). The brain’s concepts: The role of the sensory-motor system in conceptual knowledge. Cognitive Neuropsychology 22, 455–479. 10.1080/02643290442000310.

Gallivan, J.P., McLean, D.A., Smith, F.W., and Culham, J.C. (2011a). Decoding Effector-Dependent and Effector-Independent Movement Intentions from Human Parieto-Frontal Brain Activity. The Journal of Neuroscience 31, 17149–17168. 10.1523/jneurosci.1058-11.2011.

Gallivan, J.P., McLean, D.A., Valyear, K.F., and Culham, J.C. (2013). Decoding the neural mechanisms of human tool use. eLife 2, e00425. 10.7554/eLife.00425.

Gallivan, J.P., McLean, D.A., Valyear, K.F., Pettypiece, C.E., and Culham, J.C. (2011b). Decoding Action Intentions from Preparatory Brain Activity in Human Parieto-Frontal Networks. The Journal of Neuroscience 31, 9599–9610. 10.1523/jneurosci.0080-11.2011.

Garrod, S., and Pickering, M.J. (2004). Why is conversation so easy? Trends in Cognitive Sciences 8, 8–11. https://doi.org/10.1016/j.tics.2003.10.016.

Gelman, A., and Rubin, D.B. (1992). Inference from Iterative Simulation Using Multiple Sequences. Statistical Science 7, 457–472.

Geyer, S. (2004). The microstructural border between the motor and the cognitive domain in the human cerebral cortex (Springer). 10.1007/978-3-642-18910-4.

Gil-da-Costa, R., Martin, A., Lopes, M.A., Muñoz, M., Fritz, J.B., and Braun, A.R. (2006). Species-specific calls activate homologs of Broca’s and Wernicke’s areas in the macaque. Nature Neuroscience 9, 1064–1070. 10.1038/nn1741.

Gordon, R.M. (1992). The simulation theory: objections and misconceptions. Mind & Language 7, 11–34. https://doi.org/10.1111/j.1468-0017.1992.tb00195.x.

Graham, K.E., Hobaiter, C., Ounsley, J., Furuichi, T., and Byrne, R.W. (2018). Bonobo and chimpanzee gestures overlap extensively in meaning. PLOS Biology 16, e2004825. 10.1371/journal.pbio.2004825.

Hagoort, P. (2017). The core and beyond in the language-ready brain. Neuroscience & Biobehavioral Reviews 81, 194–204.

Hamilton, M.C., Schutte, N.S., and M., M.J. (1976). Hamilton anxiety scale (HAMA). In Sourcebook of Adult Assessment: Applied Clinical Psychology, (Springer), pp. 154–157.

Hanke, M., Halchenko, Y.O., Sederberg, P.B., Hanson, S.J., Haxby, J.V., and Pollmann, S. (2009). PyMVPA: a Python toolbox for multivariate pattern analysis of fMRI data. Neuroinformatics 7, 37–53. 10.1007/s12021-008-9041-y.

Hauk, O., Johnsrude, I., and Pulvermüller, F. (2004). Somatotopic representation of action words in human motor and premotor cortex. Neuron 41, 301–307. 10.1016/S0896-6273(03)00838-9.

Hellbernd, N., and Sammler, D. (2018). Neural bases of social communicative intentions in speech. Social Cognitive and Affective Neuroscience 13, 604–615. 10.1093/scan/nsy034.

Hertrich, I., Dietrich, S., and Ackermann, H. (2016). The role of the supplementary motor area for speech and language processing. Neuroscience & Biobehavioral Reviews 68, 602–610. 10.1016/j.neubiorev.2016.06.030.

Jacob, P., and Jeannerod, M. (2005). The motor theory of social cognition: a critique. Trends in Cognitive Sciences 9, 21–25. 10.1016/j.tics.2004.11.003.

Jeannerod, M. (2001). Neural simulation of action: A unifying mechanism for motor cognition. NeuroImage 14, 103–109. 10.1006/nimg.2001.0832.

Jenkinson, M., Bannister, P., Brady, M., and Smith, S. (2002). Improved Optimization for the Robust and Accurate Linear Registration and Motion Correction of Brain Images. NeuroImage 17, 825–841. https://doi.org/10.1006/nimg.2002.1132.

Jenkinson, M., Beckmann, C.F., Behrens, T.E.J., Woolrich, M.W., and Smith, S.M. (2012). FSL. NeuroImage 62, 782–790. https://doi.org/10.1016/j.neuroimage.2011.09.015.

Jenkinson, M., and Smith, S. (2001). A global optimisation method for robust affine registration of brain images. Medical Image Analysis 5, 143–156. https://doi.org/10.1016/S1361-8415(01)00036-6.

Knight, R.G. (1984). Some general population norms for the short form Beck Depression Inventory. Journal of Clinical Psychology 40, 751–753. https://doi.org/10.1002/1097-4679(198405)40:3<751::AID-JCLP2270400320>3.0.CO;2-Y.

Kriegeskorte, N., Mur, M., and Bandettini, P. (2008). Representational similarity analysis - connecting the branches of systems neuroscience. Frontiers in Systems Neuroscience 2. 10.3389/neuro.06.004.2008.

Levinson, S.C. (2016). Turn-taking in human communication – Origins and implications for language processing. Trends in Cognitive Sciences 20, 6–14. 10.1016/j.tics.2015.10.010.

Lewandowski, D., Kurowicka, D., and Joe, H. (2009). Generating random correlation matrices based on vines and extended onion method. Journal of Multivariate Analysis 100, 1989–2001. 10.1016/j.jmva.2009.04.008.

Li, X. (1983). The distribution of left and right handedness in Chinese people (中国人的左右利手分布). Acta Psychologica Sinica (心理学报) 3, 268–276.

Mayka, M.A., Corcos, D.M., Leurgans, S.E., and Vaillancourt, D.E. (2006). Three-dimensional locations and boundaries of motor and premotor cortices as defined by functional brain imaging: A meta-analysis. NeuroImage 31, 1453–1474. 10.1016/j.neuroimage.2006.02.004.

Michael, J., Sandberg, K., Skewes, J., Wolf, T., Blicher, J., Overgaard, M., and Frith, C.D. (2014). Continuous theta-burst stimulation demonstrates a causal role of premotor homunculus in action understanding. Psychological Science 25, 963–972. 10.1177/0956797613520608.

Molenberghs, P., Cunnington, R., and Mattingley, J.B. (2012). Brain regions with mirror properties: A meta-analysis of 125 human fMRI studies. Neuroscience & Biobehavioral Reviews 36, 341–349. https://doi.org/10.1016/j.neubiorev.2011.07.004.

Morgan, T.J.H., Uomini, N.T., Rendell, L.E., Chouinard-Thuly, L., Street, S.E., Lewis, H.M., Cross, C.P., Evans, C., Kearney, R., de la Torre, I., et al. (2015). Experimental evidence for the co-evolution of hominin tool-making teaching and language. Nature Communications 6, 6029. 10.1038/ncomms7029.

Oberski, D. (2014). lavaan.survey: An R Package for Complex Survey Analysis of Structural Equation Models. Journal of Statistical Software 57, 1–27. 10.18637/jss.v057.i01.

Oosterhof, N.N., Tipper, S.P., and Downing, P.E. (2012). Viewpoint (in)dependence of action representations: An MVPA study. Journal of Cognitive Neuroscience 24, 975–989. 10.1162/jocn_a_00195 %M 22264198.

Oosterhof, N.N., Tipper, S.P., and Downing, P.E. (2013). Crossmodal and action-specific: neuroimaging the human mirror neuron system. Trends in Cognitive Sciences 17, 311–318. https://doi.org/10.1016/j.tics.2013.04.012.

Pérez Hernández, L. (2001). Illocution and cognition: A constructional approach (Servicio de Publicaciones Universidad de La Rioja).

Pilgramm, S., de Haas, B., Helm, F., Zentgraf, K., Stark, R., Munzert, J., and Krüger, B. (2016). Motor imagery of hand actions: Decoding the content of motor imagery from brain activity in frontal and parietal motor areas. Human Brain Mapping 37, 81–93. 10.1002/hbm.23015.

Postle, N., McMahon, K.L., Ashton, R., Meredith, M., and de Zubicaray, G.I. (2008). Action word meaning representations in cytoarchitectonically defined primary and premotor cortices. NeuroImage 43, 634–644. https://doi.org/10.1016/j.neuroimage.2008.08.006.

Pulvermüller, F. (2005). Brain mechanisms linking language and action. Nature Reviews Neuroscience 6, 576–582. 10.1038/nrn1706.

Pulvermüller, F. (2018). Neural reuse of action perception circuits for language, concepts and communication. Progress in Neurobiology 160, 1–44. 10.1016/j.pneurobio.2017.07.001.

Pulvermüller, F., and Fadiga, L. (2010). Active perception: sensorimotor circuits as a cortical basis for language. Nature Reviews Neuroscience 11, 351–360. 10.1038/nrn2811.

Rizzolatti, G., and Arbib, M.A. (1998). Language within our grasp. Trends in Neurosciences 21, 188–194. 10.1016/S0166-2236(98)01260-0.

Rizzolatti, G., Camarda, R., Fogassi, L., Gentilucci, M., Luppino, G., and Matelli, M. (1988). Functional organization of inferior area 6 in the macaque monkey. Experimental Brain Research 71, 491–507. 10.1007/BF00248742.

Rizzolatti, G., Cattaneo, L., Fabbri-Destro, M., and Rozzi, S. (2014). Cortical Mechanisms Underlying the Organization of Goal-Directed Actions and Mirror Neuron-Based Action Understanding. Physiological Reviews 94, 655–706. 10.1152/physrev.00009.2013.

Rizzolatti, G., Fadiga, L., Gallese, V., and Fogassi, L. (1996). Premotor cortex and the recognition of motor actions. Cognitive Brain Research 3, 131–141. 10.1016/0926-6410(95)00038-0.

Rizzolatti, G., and Sinigaglia, C. (2016). The mirror mechanism: a basic principle of brain function. Nature Reviews Neuroscience 17, 757–765. 10.1038/nrn.2016.135.

Russell, B. (1950). An Inquiry into Meaning and Truth (Routledge).

Schurz, M., Radua, J., Aichhorn, M., Richlan, F., and Perner, J. (2014). Fractionating theory of mind: A meta-analysis of functional brain imaging studies. Neuroscience & Biobehavioral Reviews 42, 9–34. 10.1016/j.neubiorev.2014.01.009.

Searle, J.R. (1969). Speech acts: An essay in the philosophy of language (Cambridge University Press).

Searle, J.R. (1985). Expression and meaning: Studies in the theory of speech acts (Cambridge University Press).

Searle, J.R., and Vanderveken, D. (1985). Foundations of illocutionary logic (Cambridge University Press).

Seed, A., and Byrne, R. (2010). Animal tool-use. Current Biology 20, R1032–R1039. https://doi.org/10.1016/j.cub.2010.09.042.

Seyfarth, R.M., Cheney, D.L., and Marler, P. (1980). Monkey responses to three different alarm calls: Evidence of predator classification and semantic communication. Science 210, 801–803. doi:10.1126/science.7433999.

Shibata, M., Abe, J.-i., Itoh, H., Shimada, K., and Umeda, S. (2011). Neural processing associated with comprehension of an indirect reply during a scenario reading task. Neuropsychologia 49, 3542–3550. 10.1016/j.neuropsychologia.2011.09.006.

Smith, S.M. (2002). Fast robust automated brain extraction. Human Brain Mapping 17, 143–155. https://doi.org/10.1002/hbm.10062.

Stelzer, J., Chen, Y., and Turner, R. (2013). Statistical inference and multiple testing correction in classification-based multi-voxel pattern analysis (MVPA): Random permutations and cluster size control. NeuroImage 65, 69–82. 10.1016/j.neuroimage.2012.09.063.

Stout, D., and Chaminade, T. (2012). Stone tools, language and the brain in human evolution. Philosophical Transactions of the Royal Society B: Biological Sciences 367, 75–87. 10.1098/rstb.2011.0099.

Stout, D., Toth, N., Schick, K., and Chaminade, T. (2008). Neural correlates of early stone age toolmaking: Technology, language and cognition in human evolution. Philosophical Transactions of the Royal Society B: Biological Sciences 363, 1939–1949. 10.1098/rstb.2008.0001.

Thibault, S., Py, R., Gervasi, A.M., Salemme, R., Koun, E., Lövden, M., Boulenger, V., Roy, A.C., and Brozzoli, C. (2021). Tool use and language share syntactic processes and neural patterns in the basal ganglia. Science 374, eabe0874. doi:10.1126/science.abe0874.

Thompson, J.A., Basista, M.J., Wu, W., Bertram, R., and Johnson, F. (2011). Dual Pre-Motor Contribution to Songbird Syllable Variation. The Journal of Neuroscience 31, 322–330. 10.1523/jneurosci.5967-09.2011.

Tylén, K., Weed, E., Wallentin, M., Roepstorff, A., and Frith, C.D. (2010). Language as a Tool for Interacting Minds. Mind & Language 25, 3–29. 10.1111/j.1468-0017.2009.01379.x.

van Ackeren, M.J., Casasanto, D., Bekkering, H., Hagoort, P., and Rueschemeyer, S.-A. (2012). Pragmatics in action: Indirect requests engage theory of mind areas and the cortical motor network. Journal of Cognitive Neuroscience 24, 2237–2247. 10.1162/jocn_a_00274 %M 22849399.

Watson, Stuart K., Townsend, Simon W., Schel, Anne M., Wilke, C., Wallace, Emma K., Cheng, L., West, V., and Slocombe, Katie E. (2015). Vocal learning in the functionally referential food grunts of chimpanzees. Current Biology 25, 495–499. https://doi.org/10.1016/j.cub.2014.12.032.

Willems, R.M., Labruna, L., D’Esposito, M., Ivry, R., and Casasanto, D. (2011). A functional role for the motor system in language understanding: Evidence from theta-burst transcranial magnetic stimulation. Psychological Science 22, 849–854. 10.1177/0956797611412387.

Wilson, S.M., Saygin, A.P., Sereno, M.I., and Iacoboni, M. (2004). Listening to speech activates motor areas involved in speech production. Nature Neuroscience 7, 701–702. 10.1038/nn1263.

Wittgenstein, L. (1953). Philosophical investigations (Blackwell).

Zhang, Y., Brady, M., and S., S. (2001). Segmentation of brain MR images through a hidden Markov random field model and the expectation-maximization algorithm. IEEE Transactions on Medical Imaging 20, 45–57. 10.1109/42.906424.

Zwaan, R.A. (2016). Situation models, mental simulations, and abstract concepts in discourse comprehension. Psychonomic Bulletin & Review 23, 1028–1034. 10.3758/s13423-015-0864-x.

