## Supplemental Information for "Representing linguistic communicative goals in the premotor cortex"

### Supplemental Methods

*Pilot evaluation of the scripts for the fMRI study*

Prior to the fMRI experiment, we created 100 quadruplets of scripts and conducted a pilot evaluation (Pilot 1) to assess the acceptability of the predefined communicative goals conveyed by critical sentences and the extent to which communicative goals can be predicted by the ten features based on Pérez Hernández's corpus study (2001), including speaker's will, cost-benefit, and pleasure[i], addressee's will, cost-benefit, and pleasure, social distance, relative power, and mitigation. Forty-eight native Chinese speakers (35 females, mean age = 22 years, *SD* = 3, *range*: [17, 29]) without a known history of psychiatric or neurological disorders participated in Pilot 1. None of them participated in the fMRI experiment or the lesion study.

The 100 quadruplets of scripts were divided into four lists according to a Latin-square procedure. Firstly, to assess the acceptability of the predefined communicative goal of the critical sentence of each script, participants were asked to read the script and complete a forced-choice question with 7 alternatives. The alternatives included two assertives (to reply and to state), two commissives (to promise and to assure), two

---

[i] Although Pérez Hernández (2001) did not consider the interlocutors' pleasure as a feature of linguistic communication, we nevertheless included it in the rating because a previous study have suggested the potential impact of emotional information on language processes (Havas et al., 2007).

directives (to request and to order), and “other”. Participants were instructed to choose the most appropriate option for each described scenario, and to choose “other” only if all of the six alternatives of communicative goals were considered inappropriate. Because in the current context the two alternatives attributed to the same category can be used to describe the same communicative goal, the predefined communicative goal was considered as accepted by the participants when either of them was chosen.

Secondly, participants were asked to rate each of the ten features for each script. A 7-point scale was used for each of these features: (1) performer’s capability, from *1* (the performer is totally not capable of conducting the task described by the critical sentence) to *7* (the performer is very capable of conducting the task); (2) speaker’s will and (3) addressee’s will, from *1* (the speaker/addressee is very unwilling to conduct the task) to *7* (the speaker/addressee is very willing to conduct the task); (4) speaker’s cost-benefit and (5) addressee’s cost-benefit, from *1* (the speaker/addressee would take a high cost if the performer has accomplished the task) to *7* (the speaker/addressee would be benefitted highly if the performer has accomplished the task); (6) speaker’s pleasure and (7) addressee’s pleasure, from *1* (the speaker/addressee is very unpleased when communicating) to *7* (the speaker/addressee is very pleased when communicating); (8) relative power, from *1* (the addressee’s power is definitely higher than the speaker’s) to *7* (the speaker’s power is definitely higher than the addressee's); (9) social distance between the interlocutors, from *1* (very close) to *7* (very remote); and (10) the mitigation of the critical sentence, from *1* (not mitigated at all) to *7* (highly mitigated).

To examine the extent to which communicative goals could be predicted by the ten features, the same Bayesian logistic mixed modelling as the analyses of the post-scanning ratings in the fMRI experiment was conducted on the 80 scripts selected based on the results of Pilot 1 (see below).

To investigate the potential accounting factor(s) for the rating of the ten features, exploratory factor analysis (EFA) was performed on the ratings of the 80 scripts using the *psych* package (Revelle, 2017) in R. To decide the number of factors in a model, parallel analysis with 50,000 iterations was performed using the *fa.parallel* function, resulting in a solution of a three-factor model. Principle axis factor analysis with three factors based on oblique rotation was then implemented using the *fa* function.

*ROI-based univariate analysis of the fMRI data.*

To examine univariate activities in each of the ROIs, parameter estimates of the critical sentences were extracted from the individual-level GLMs (see *Methods*) using *Featquery* (http://poc.vl-e.nl/distribution/manual/fsl-3.2/feat5/featquery.html). Differences in parameter estimates were tested by paired-sample *t*-test for the four pair-wise comparisons: (1) *Promise* vs. *Reply-1*; (2) *Request* vs. *Reply-2*; (3) *Promise* vs. *Request*; (4) *Reply-1* vs. *Reply-2*. Hence, 24 (6 ROIs × 4 pair-wise comparisons) comparisons were conducted. Statistical significance was determined by a Bonferroni corrected threshold of $p < 0.002$ (= 0.05/24).

*Multivariate pattern classifications (MVPCs) for medial prefrontal cortex and left/right temporo-parietal junction*

As previous studies have shown the activations of typical "theory of mind" regions, including the medial prefrontal cortex (MPFC) and temporo-parietal junction (TPJ), in linguistic communication tasks (Feng et al., 2017; Feng et al., 2021; Shibata et al., 2011), we conducted additional MVPCs for these regions. We defined ROIs for the MPFC, left TPJ, and right TPJ (*Figure S4a*) based on the *NeuroSynth* meta-analytic database (Yarkoni et al., 2011). We firstly obtained the activation map related to "theory of mind" by searching the term "theory mind" on the *NeuroSynth* platform (https://neurosynth.org/) and secondly extracted the three ROIs from this activation map. To test whether the ROIs represent the information on communicative goals and whether the two premotor ROIs represent more information relative to these ROIs, ROI-based MVPCs and combinatorial MVPCs were conducted for these ROIs with the same methods used for the premotor ROIs and the perisylvian ROIs. For permutation-based significance tests with Bonferroni correction for multiple comparisons, classification accuracy in each of the three ROIs for each pair-wise classification was tested with a threshold of $p < 0.004$ (= 0.05/12) as there were 12 (3 “theory of mind” ROIs × 4 pair-wise classifications) comparisons, each improvement in accuracy contributed by an added ROI for an initial ROI was tested with a threshold of $p < 0.002$ (= 0.05/24) as there were 24 (2 premotor ROIs × 3 “theory of mind” ROIs × 2 pair-wise classifications × 2 alternatives of initial-added ROIs pair) comparisons, and each difference between the improvement in accuracy contributed by a premotor ROI for either of the MPFC, left TPJ, or right TPJ and the improvement contributed by either of the MPFC, left TPJ, or right TPJ for a premotor ROI was tested with a threshold of $p < 0.004$ (= 0.05/12) as there were 12 (2 premotor ROIs × 3 “theory of mind” ROIs × 2 pair-wise classifications) comparisons.

*Methods of the pilot studies for the lesion study*

To accommodate the patients’ cognitive states, prior to the lesion study, we adopted and simplified the scripts used in the fMRI experiment while keeping the structure of these scripts unchanged, to create another 100 quadruplets of scripts. The pilot evaluation of these scripts (Pilot 2) was conducted with 92 native Chinese speakers (65 females, mean age = 21 years, *SD* = 3, *range*: [18, 33]) without a known history of psychiatric or neurological disorders. This larger sample size, compared to the size of the Pilot 1, was used to replicate the previous results with a higher reliability. The procedure and the methods of data analyses were the same as those used in the Pilot 1.

Moreover, to assess the appropriateness of the behavioral experimental procedure for the lesion study, a pilot experiment (Pilot 3) was conducted with another group of 40 native Chinese speakers (28 females, mean age = 22 years, *SD* = 2, *range*: [18, 27]) without a known history of psychiatric or neurological disorders. The experimental procedure was the same as that in the lesion study, but with a shorter time limit for the presentations of the scripts and for the responses (*Figure S1*). None of the participants in these two pilot studies participated in the main lesion study or the fMRI study.

The Bayesian hierarchical logistic models were fitted to assess the extent to which communicative goals could be predicted by the ratings of the speaker's will and the addressee's will for "*Promise* vs. *Reply-1*" and "*Request* vs. *Reply-2*" respectively. The model parameters were estimated at both the group-level and the participant-level. The settings of variables and priors and the model-fitting method were the same as in the main experiment (see *Methods*). The model-fitting only included trials with correct performer judgements (93% of the data).

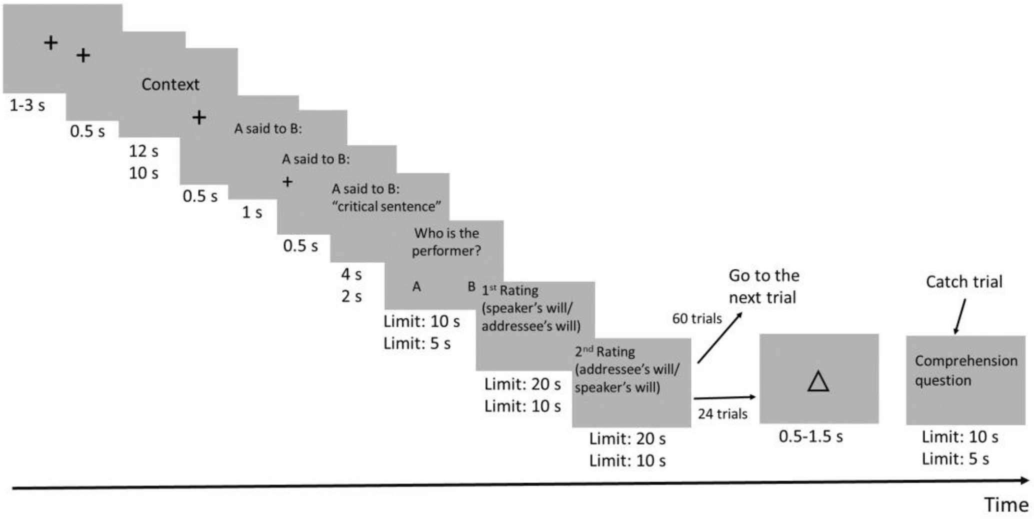


**Figure S1 |** Procedure of the lesion study and the Pilot 3. The durations of the presentation of scripts and the time limit for responses were written in two rows, for illustration purposes. Each duration/ time limit written in the first row was for the main lesion study, while the one written in the second row was for Pilot 3.

### Supplemental Results

*Pilot evaluation of the scripts of the fMRI study*

For the initially created 100 quadruplets of scripts, we evaluated their appropriateness for the experimental design by considering both the results of Pilot 1 (see below) and the content of these scripts. Eighty quadruplets of scripts were selected and further used in the fMRI experiment. Here we report the pilot evaluative results of the 80 selected quadruplets of scripts. The predefined communicative goals for these scripts had mean acceptance rates of 90% or higher (*Table S1*).

**Table S1 |** Acceptability of the predefined communicative goals in different scripts

| | Communicative goal | *Mean* | *SD* | Range |
|---|---|---|---|---|
| **Pilot1: pilot evaluation for the fMRI study** | *Promise* | 94% | 6% | [83%, 100%] |
| | *Reply-1* | 95% | 6% | [75%, 100%] |
| | *Request* | 94% | 6% | [83%, 100%] |
| | *Reply-2* | 90% | 9% | [75%, 100%] |
| **Pilot 2: pilot evaluation for the lesion study** | *Promise* | 93% | 7% | [70%, 96%] |
| | *Reply-1* | 90% | 6% | [74%, 100%] |
| | *Request* | 84% | 8% | [70%, 100%] |
| | *Reply-2* | 83% | 7% | [70%, 100%] |

*SD*, standard deviation

The Bayesian logistic mixed modelling obtained the following results for the 80 selected scripts (*Figure S2a* and *Table S2*). (1) "*Promise* vs. *Reply-1*" was predicted by ratings of addressee's will, addressee's cost-benefit, addressee's pleasure, speaker's cost-benefit, performer's capability and social distance, with higher ratings of addressee's will, cost-benefit, and pleasure and social distance, and lower ratings of speaker's cost-benefit for *Promise* than for *Reply-1*. (2) "*Request* vs. *Reply-2*" was predicted by ratings of speaker's will, speaker's cost-benefit, speaker's pleasure, addressee's cost-benefit and the mitigation, with higher ratings of speaker's will, cost-benefit and pleasure, and lower ratings of addressee's cost-benefit and mitigation for *Request* than for *Reply-2*.

The EFA showed that the ratings of the features were explained by three factors (*Figure S2b*): (1) speaker's attitudes, explaining 48% of the variance, with the largest loadings on speaker's will (0.75), speaker's cost-benefit (0.72), and speaker's pleasure (0.52); (2) addressee's attitudes, explaining 29% of the variance, with the largest loadings on addressee's will (0.97), addressee's cost-benefit (0.82), and addressee's pleasure (0.88); (3) contextual information, explaining 22% of the variance, with the largest loadings on social distance (-0.55), relative power (0.44), and mitigation (0.73). The ratings of performer's capability had absolute loadings lower than 0.3 and were not considered as explainable by either factor.

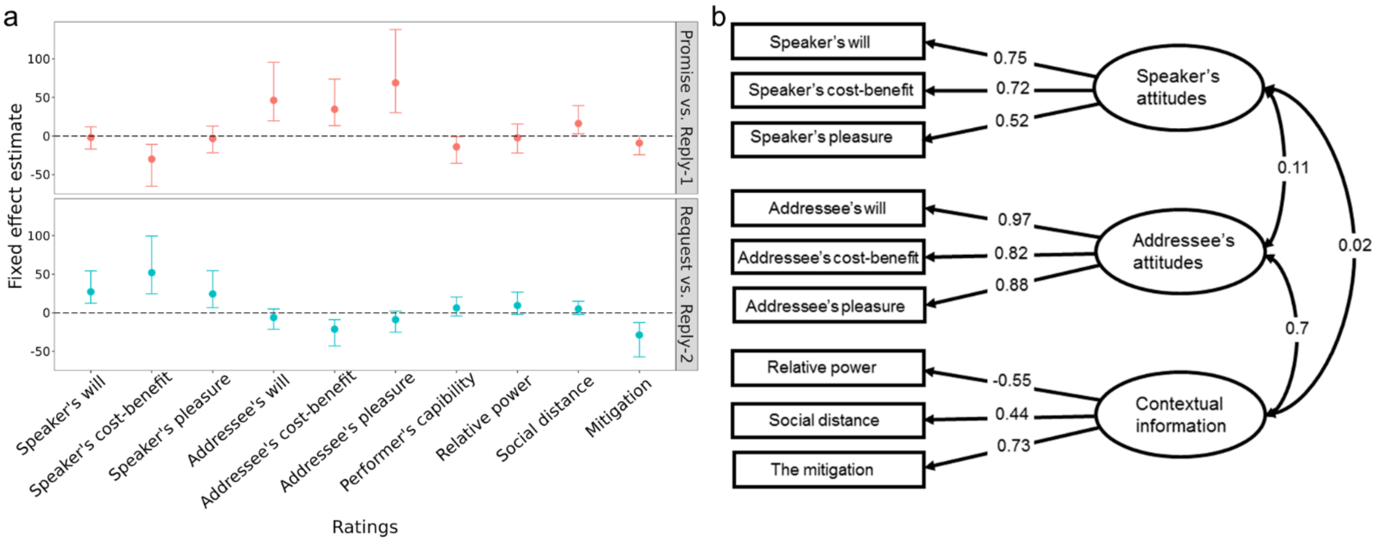


**Figure S2 | Results of Pilot 1 which evaluated the scripts for the fMRI study. a,** Results of Bayesian logistic mixed models of the ratings. The posterior estimate of the fixed effect (vertical axis) for each contextual feature (horizontal axis) in the "*Promise* vs. *Reply-1*" model (the upper panel, red) and the "*Request* vs. *Reply-2*" model (the lower panel, turquoise) are plotted. **b,** EFA result of the feature ratings. The ellipses represent the accounting factors and the rectangles represent the contextual features. The correlations between the accounting factors and the loadings of the accounting factors on the contextual features are embedded in the arrows.

**Table S2 | Results of Bayesian logistic mixed models for the ratings**

| Model | Contextual feature | The fMRI study | | | | The lesion study | |
|---|---|---|---|---|---|---|---|
| | | Pilot evaluation | | Post-scanning sheets | | Pilot study 1 | |
| | | *b* | 95% *CrI* | *b* | 95% *CrI* | *b* | 95% *CrI* |
| *Promise* | Speaker's will | -1.85 | [-16.92, 11.9] | -0.1 | [-0.75, 0.44] | **-0.47** | **[-0.78, -0.19]** |
| vs. | Speaker's cost-benefit | **-29.88** | **[-64.98, -10.8]** | **-1.18** | **[-2.04, -0.48]** | **-0.75** | **[-1.15, -0.41]** |
| *Reply-1* | Speaker's pleasure | -3.17 | [-21.65, 13.03] | -0.35 | [-1.19, 0.35] | 0.34 | [-0.07, 0.77] |
| | Addressee's will | **46.32** | **[19.78, 95.38]** | **2.21** | **[1.52, 3.09]** | **1.62** | **[1.25, 2.05]** |
| | Addressee's cost-benefit | **34.67** | **[13.48, 73.73]** | **1.77** | **[1.04, 2.71]** | **1.46** | **[1.07, 1.92]** |
| | Addressee's pleasure | **68.93** | **[30.33, 138]** | **3.28** | **[2.32, 4.55]** | **2.05** | **[1.63, 2.54]** |
| | Performer's capability | **-13.99** | **[-35.25, -0.81]** | -0.5 | [-1.16, 0.11] | **-0.48** | **[-0.8, -0.21]** |
| | Relative power | -2.35 | [-21.87, 15.67] | -1 | [-2.2, 0.03] | 0.44 | [-0.05, 0.96] |
| | Social distance | **16.23** | **[2.91, 39.32]** | 0.21 | [-0.26, 0.73] | **0.57** | **[0.32, 0.83]** |
| | The mitigation | -9.06 | [-24.3, 0.28] | -0.06 | [-0.64, 0.5] | **-0.32** | **[-0.58, -0.08]** |
| *Request* | Speaker's will | **27.21** | **[12.32, 54.19]** | **1.47** | **[1.05, 2]** | **1.26** | **[0.96, 1.59]** |
| vs. | Speaker's cost-benefit | **52.01** | **[24.69, 99.44]** | **2.35** | **[1.7, 3.15]** | **2.04** | **[1.62, 2.51]** |
| *Reply-2* | Speaker's pleasure | **24.42** | **[6.75, 54.45]** | **1.00** | **[0.3, 1.85]** | **0.81** | **[0.31, 1.35]** |
| | Addressee's will | -6.11 | [-21.27, 5.04] | **-0.60** | **[-0.99, -0.25]** | **-0.62** | **[-0.94, -0.34]** |
| | Addressee's cost-benefit | **-21.24** | **[-43.14, -8.91]** | **-0.57** | **[-1.02, -0.19]** | **-0.83** | **[-1.2, -0.49]** |
| | Addressee's pleasure | -8.90 | [-25.17, 2.02] | **-0.52** | **[-1.05, -0.06]** | **-1.13** | **[-1.57, -0.75]** |
| | Performer's capability | 6.34 | [-4.05, 20.55] | -0.05 | [-0.41, 0.32] | -0.21 | [-0.47, 0.03] |
| | Relative power | 9.39 | [-2.14, 26.65] | **0.67** | **[0.11, 1.28]** | 0.20 | [-0.2, 0.61] |
| | Social distance | 4.92 | [-2.26, 14.99] | 0.16 | [-0.18, 0.5] | **0.37** | **[0.14, 0.61]** |
| | The mitigation | **-28.72** | **[-57.12, -12.67]** | **-1.03** | **[-1.48, -0.65]** | **-1.05** | **[-1.39, -0.75]** |

*b*, mean posterior estimate; 95% *CrI*, 95% credible interval

Bold texts denote that the 95% *Crl* excluding 0, suggesting statistically significant effects.

*ROI-based univariate analyses*

Results of the fMRI univariate analyses showed that activations in LMTG were higher for *Request* than for either *Promise* or *Reply-2* (*Figure S3*; $t_{(55)}$ = 4.2, $p$ = 0.0001; $t_{(55)}$ = 3.58, $p$ = 0.0007, respectively). No other significant difference was observed.

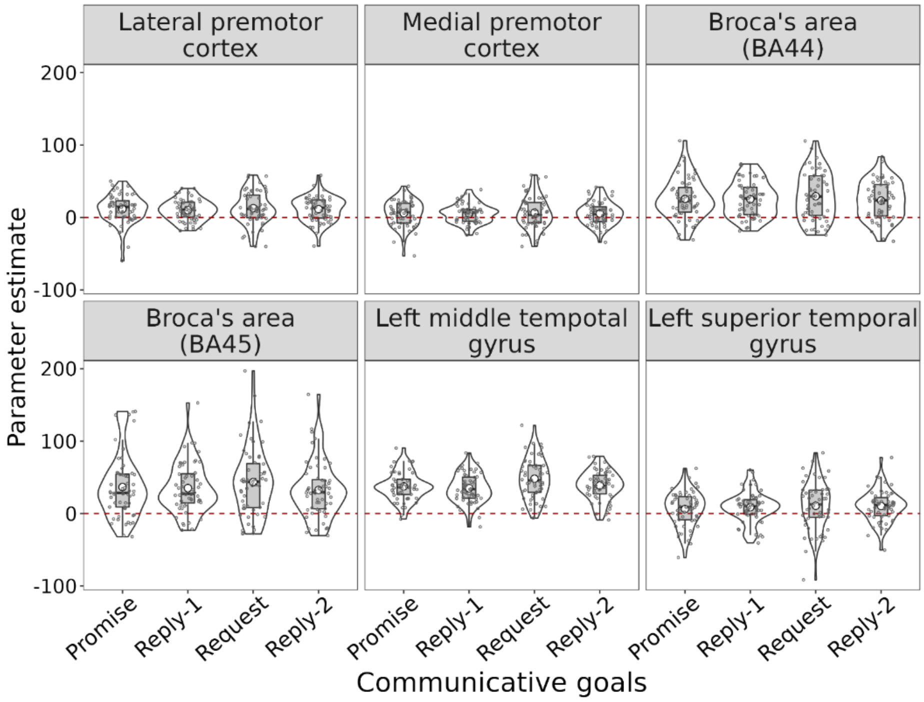


**Figure S3 | Results of ROI-based univariate analyses in the fMRI study.**

*MVPCs for MPFC and left/right TPJ*

Results of ROI-based MVPCs (*Figure S4b* and *Table S3*) showed that, for "*Promise* vs. *Reply-1*", classification accuracies were above chance level in the left TPJ and right TPJ (all *p*-values < .0005). For "*Request* vs. *Reply-2*", accuracies were above chance level in all the three ROIs (all *p*-values < .0005). For "*Promise* vs. *Request*", accuracies were above chance level in the MPFC and right TPJ (all *p*-values < .0005). For "*Reply-1* vs. *Reply-2*", no significant effect was observed.

For results of combinatorial MVPCs (*Figure S4c* and *Table S3*), using either of the MPFC, left TPJ, or right TPJ as the initial ROI and LPMC or MPMC as the added ROI, in classifying either "*Promise* vs. *Reply-1*" or "*Request* vs. *Reply-2*", both LPMC and MPMC significantly improved classification accuracy in either MPFC, left TPJ, or right TPJ (all *p*-values < 0.0005). In contrast, using LPMC or MPMC as the initial ROI and either of MPFC, left TPJ, or right TPJ as the added ROI, only the left TPJ improved the accuracy in MPMC in classifying "*Request* vs. *Reply-2*" ($p <$ 0.0005).

For results of the comparisons between the improvements in classification accuracy contributed by LPMC or MPMC for either MPFC, left TPJ, or right TPJ and that contributed by either MPFC, left TPJ, or right TPJ for LPMC or MPMC, in classifying either "*Promise* vs. *Reply-1*" or "*Request* vs. *Reply-2*", the improvement contributed by either premotor ROI was significantly higher than the improvement contributed by either MPFC, left TPJ, or right TPJ (all *p*-values < 0.0005).

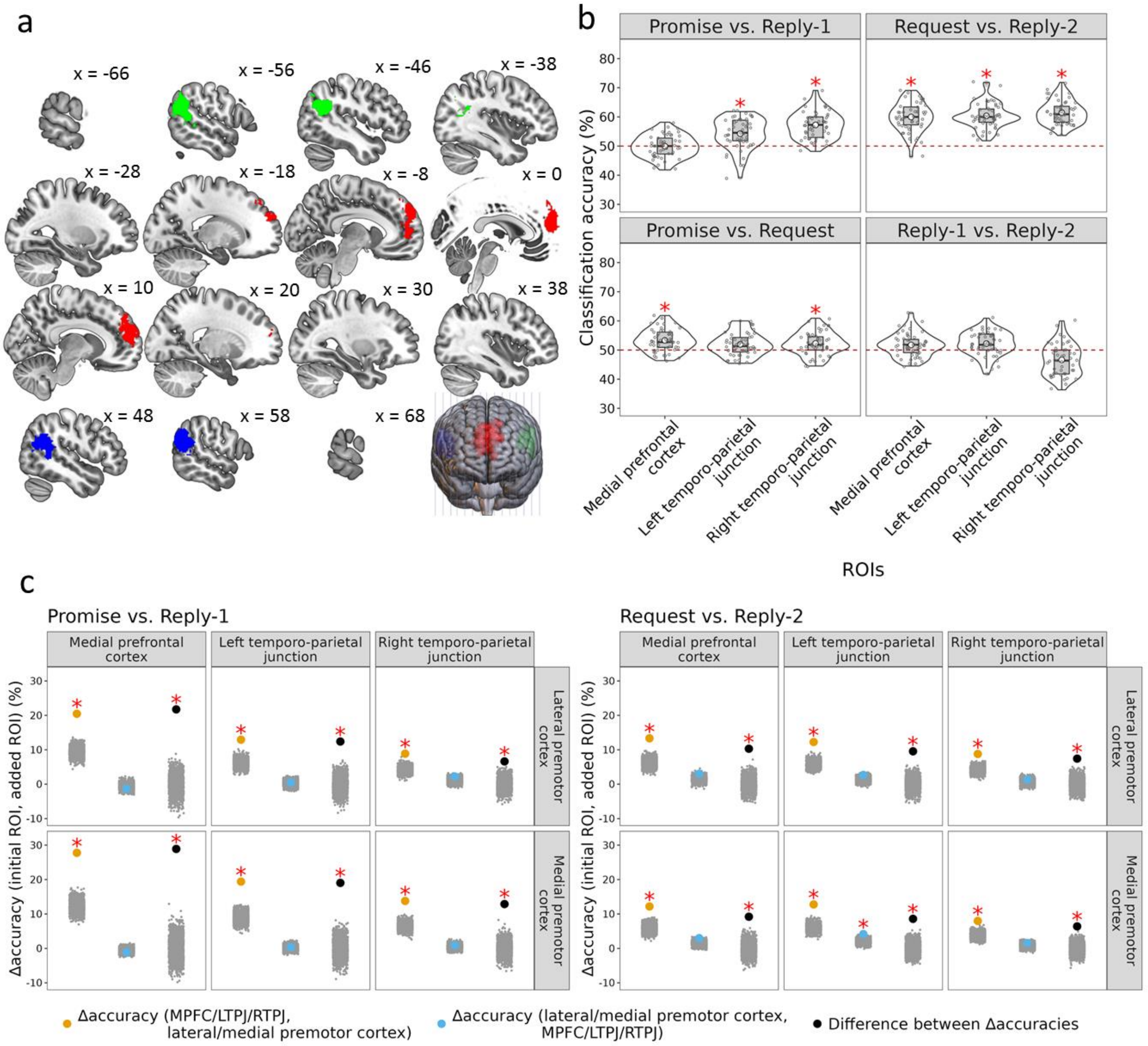


**Figure S4 | Results of additional MVPCs. a,** Three "theory of mind" ROIs were defined based on the NeuroSynth meta-analytic database. Red, medial prefrontal cortex; green, left temporo-parietal junction; blue, right temporo-parietal junction (x-coordinates based on the MNI system). **b,** Results of ROI-based MVPCs. The classification accuracies (vertical axis) in the ROIs (horizontal axis) for the four pair-wise classifications are illustrated. Left top, *Promise* vs. *Reply-1*; right top, *Request* vs. *Reply-2*; left bottom, *Promise* vs. *Request*; right bottom, *Reply-1* vs. *Reply-2*. The red dashed lines represent the chance-level percentage of binary classification (50%). Red stars represent the statistical significance of permutation tests with Bonferroni correction. **c,** Results of combinatorial MVPCs. Left panel, *Promise* vs. *Reply-1*; right panel, *Request* vs. *Reply-2*. Vertical axes illustrate the improvement in classification accuracy contributed by an added ROI for an initial ROI. Red stars represent the statistical significance of permutation tests. For abbreviations in the legend, MPFC, medial prefrontal cortex; LTPJ, left temporo-parietal junction; RTPJ, right temporo-parietal junction. Each yellow dot indicates the improvement in classification accuracy contributed by a premotor ROI for either MPMC, LTPJ, or RTPJ. Each blue dot indicates the improvement in classification accuracy contributed by either MPMC, LTPJ, or RTPJ for a premotor ROI. Each black dot indicates the difference between the improvement in classification accuracy contributed by a premotor ROI for either MPMC, LTPJ, or RTPJ and that contributed by either MPMC, LTPJ, or RTPJ for a premotor ROI. The crowded small gray dots indicate data points of null distributions for permutation tests.

### Table S3 | Results of additional MVPCs

ROI-based MVPCs

| ROI (n volxels) | Index | Pair-wise classification | | | |
|---|---|---|---|---|---|
| | | *Promise* vs. *Reply-1* | *Request* vs. *Reply-2* | *Promise* vs. *Request* | *Reply-1* vs. *Reply-2* |
| MPFC (1962) | *Accuracy* (%) | **50** | **60** | **53** | 52 |
| | *p* | **0.639** | **< 0.0005** | **< 0.0005** | 0.025 |
| LTPJ (1308**)** | *Accuracy* (%) | **54** | **60** | 52 | 52 |
| | *p* | **< 0.0005** | **< 0.0005** | 0.01 | 0.011 |
| RTPJ (1519) | *Accuracy* (%) | **57** | **61** | **52** | 47 |
| | *p* | **< 0.0005** | **< 0.0005** | **< 0.0005** | > 0.999 |

Combinatorial MVPCs of Δ*accuracy (initial ROI, added ROI)*

| Pair-wise classification | Premotor ROI | Perisylvian ROI | Δ*accuracy (perisylvian ROI, premotor ROI)* | | Δ*accuracy (premotor ROI, perisylvian ROI)* | | Difference between Δ*accuracies* | |
|---|---|---|---|---|---|---|---|---|
| | | | Δ*accuracy* (%) | *p* | Δ*accuracy* (%) | *p* | Δ*accuracy* (%) | *p* |
| *Promise* vs. *Reply-1* | LPMC | MPFC | **20** | **< 0.0005** | -1 | 0.2 | **22** | **< 0.0005** |
| | | LTPJ | **13** | **0.0015** | 1 | 0.496 | **12** | **< 0.0005** |
| | | RTPJ | **9** | **< 0.0005** | 2 | 0.047 | **7** | **< 0.0005** |
| | MPMC | MPFC | **28** | **< 0.0005** | -1 | 0.183 | **29** | **< 0.0005** |
| | | LTPJ | **19** | **< 0.0005** | 0.3 | 0.577 | **19** | **< 0.0005** |
| | | RTPJ | **14** | **< 0.0005** | 0.9 | 0.235 | **13** | **< 0.0005** |
| *Request* vs. *Reply-2* | LPMC | MPFC | **13** | **< 0.0005** | 3 | 0.01 | **10** | **< 0.0005** |
| | | LTPJ | **12** | **< 0.0005** | 3 | 0.012 | **10** | **< 0.0005** |
| | | RTPJ | **9** | **< 0.0005** | 1 | 0.016 | **7** | **< 0.0005** |
| | MPMC | MPFC | **12** | **< 0.0005** | 3 | 0.005 | **9** | **< 0.0005** |
| | | LTPJ | **13** | **< 0.0005** | **4** | **< 0.0005** | **9** | **< 0.0005** |
| | | RTPJ | **8** | **< 0.0005** | 2 | 0.104 | **6** | **< 0.0005** |

Texts in bold indicate statistical significance for the permutation-based significance testing with Bonferroni correction. For ROI-based MVPCs, accuracies were tested with a Bonferroni corrected significance threshold of $p < 0.004$. For combinatorial MVPCs, statistical significances for the permutation tests at the first step and the second step were determined by Bonferroni corrected thresholds of $p < 0.002$ and $p < 0.004$ respectively. MPFC, medial prefrontal cortex; LTPJ, left temporo-parietal junction; RTPJ, right temporo-parietal junction.

*Pilot studies for the lesion study*

In Pilot 2, the predefined communicative goal of each of the 84 selected scripts (out of 100) had an acceptance rate of 83% or higher (*Table S1*).

Bayesian logistic mixed modelling showed that, in the "*Promise* vs. *Reply-1*" model, the ratings of addressee's will, cost-benefit, and pleasure and social distance were significantly higher for *Promise* than for *Reply-1*, whereas the ratings of speaker's will and cost-benefit, performer's capability, and mitigation were lower for *Promise* than for *Reply-1*(*Figure S5a* and *Table S2*). In the "*Request* vs. *Reply-2*" model, the ratings of speaker's will, cost-benefit, and pleasure and social distance were higher for *Request* than for *Reply-2*, whereas the ratings of addressee's will, cost-benefit, and pleasure and the mitigation were lower for *Request* than for *Reply-2*.

In Pilot 3, the mean accuracy of the performer judgement was 93% (*SD* = 9%), the mean accuracy of the responses to the comprehension questions was 84% (*SD* = 8%). Bayesian hierarchical logistic modelling showed that: (1) the addressee's will ratings were higher for *Promise* than for *Reply-1*; (2) the speaker's will ratings were higher for *Request* than for *Reply-2,* whereas the addressee's will ratings had a reversed pattern (*Figure S5b* and *Table S4*).

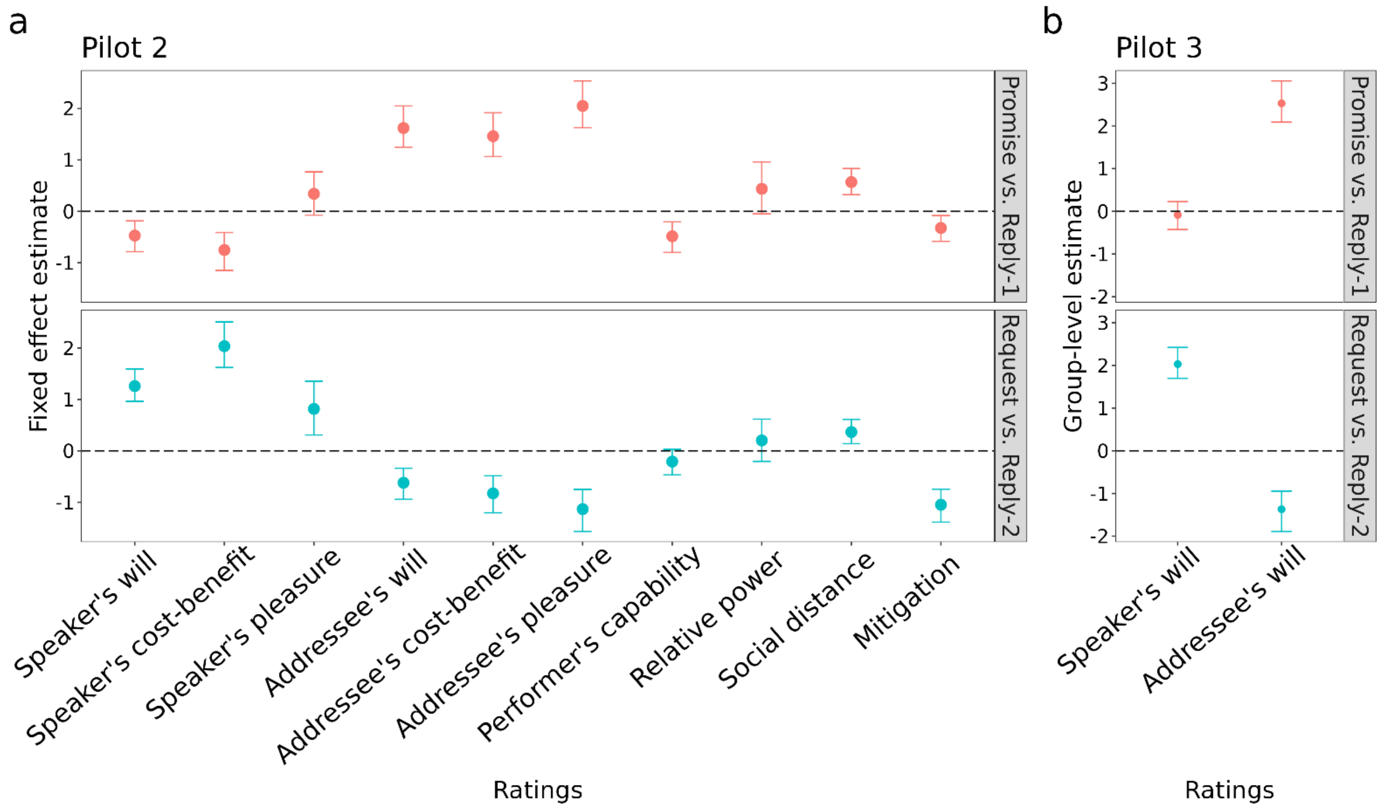


**Figure S5 | Results of the pilot studies for the lesion study. a,** Results of Bayesian logistic mixed modelling for Pilot 2 which evaluated the scripts for the lesion study. The posterior estimate of the fixed slope (vertical axis) for each rating of the feature (horizontal axis) is plotted. **b,** Results of Bayesian hierarchical logistic modelling for Pilot 3. The posterior estimates of the ratings (vertical axis) of the speaker's will and the addressee's will (horizontal axis) are plotted. The upper panel represents the "*Promise* vs. *Reply-1*" model (red), the lower panel represents the "*Request* vs. *Reply-2*" model (turquoise). The solid circles represent mean posterior estimates. The error bars represent 95% *CrI*.

**Table S4 | Results of Bayesian hierarchical logistic modelling in the lesion study**

| Model | Predictor | Patients with premotor lesions | | Lesion controls | | Healthy controls | | Pilot 3 | |
|---|---|---|---|---|---|---|---|---|---|
| | | *b* | 95% *CrI* | *b* | 95% *CrI* | *b* | 95% *CrI* | *b* | 95% *CrI* |
| *Promise* vs. | Speaker's will | -0.26 | [-0.61,0.08] | -0.0008 | [-0.41,0.4] | 0.21 | [-0.1,0.51] | -0.08 | [-0.43, 0.23] |
| *Reply-1* | Addressee's will | 0.68 | [-0.05,1.43] | **1.19** | **[0.43,1.99]** | **1.89** | **[1.33,2.5]** | **2.53** | **[2.09, 3.06]** |
| *Request* vs. | Speaker's will | 0.44 | [-0.21,1.09] | **0.85** | **[0.17,1.59]** | **1.38** | **[0.89,1.92]** | **2.03** | **[1.7, 2.42]** |
| *Reply-2* | Addressee's will | **-1.38** | **[-2.07,-0.74]** | **-0.99** | **[-1.7,-0.32]** | **-1.28** | **[-1.89,-0.76]** | **-1.37** | **[-1.89, -0.95]** |

*b*, mean posterior estimate; 95% *CrI*, 95% credible interval.

Bold texts denote that the 95% *Crl* excluding 0, suggesting statistically significant effects.

### References

Feng, W., Wu, Y., Jan, C., Yu, H., Jiang, X., and Zhou, X. (2017). Effects of contextual relevance on pragmatic inference during conversation: An fMRI study. Brain and Language *171*, 52-61. 10.1016/j.bandl.2017.04.005.

Feng, W., Yu, H., and Zhou, X. (2021). Understanding particularized and generalized conversational implicatures: Is theory-of-mind necessary? Brain and Language *212*, 104878. https://doi.org/10.1016/j.bandl.2020.104878.

Havas, D.A., Glenberg, A.M., and Rinck, M. (2007). Emotion simulation during language comprehension. Psychonomic Bulletin & Review *14*, 436-441. 10.3758/BF03194085.

Pérez Hernández, L. (2001). Illocution and cognition: A constructional approach (Servicio de Publicaciones Universidad de La Rioja).

Revelle, W. (2017). psych: Procedures for Personality and Psychological Research. Software.

Shibata, M., Abe, J.-i., Itoh, H., Shimada, K., and Umeda, S. (2011). Neural processing associated with comprehension of an indirect reply during a scenario reading task. Neuropsychologia *49*, 3542-3550. 10.1016/j.neuropsychologia.2011.09.006.

Yarkoni, T., Poldrack, R.A., Nichols, T.E., Van Essen, D.C., and Wager, T.D. (2011). Large-scale automated synthesis of human functional neuroimaging data. Nature Methods *8*, 665-670. 10.1038/nmeth.1635.

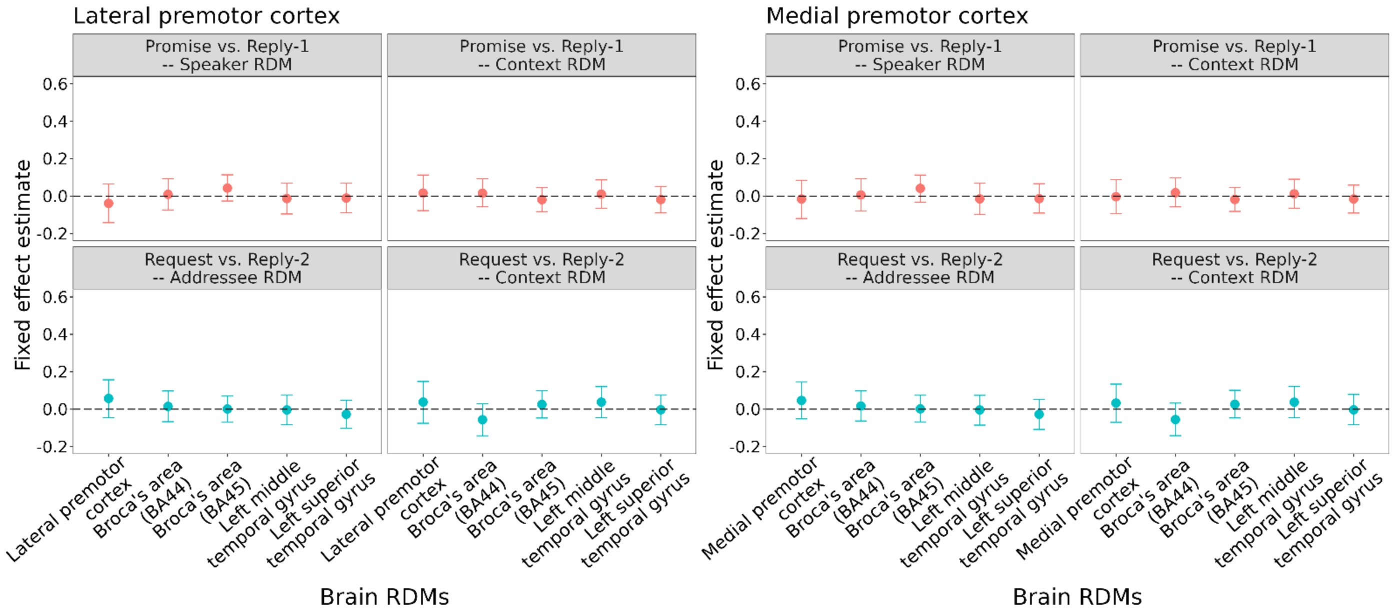


**Figure S6 | Supplemental results the RS decoding models.** Upper panel illustrates the models with Speaker RDM and Context RDM as response variables for "*Promise* vs. *Reply-1*" (red); lower panels illustrates the models with Addressee RDM and Context RDM as response variables for "*Request* vs. *Reply-2*" (turquoise). Left panel illustrates the models with the LPMC RDM and the RDMs of the prerisylvian ROIs as predictors; right panel illustrates the models with the MPMC RDM and the RDMs of the perisylvian ROIs as predictors.

### Table S5 | Supplemental results of RS decoding models

**Representational similarity decoding model**

| | | | ROI | | | | | | | | | |
|---|---|---|---|---|---|---|---|---|---|---|---|---|
| | | | LPMC/MPMC | | Left BA44 | | Left BA45 | | LMTG | | LSTG | |
| Model | Prediction | Behavioral RDM | *b* | 99.92% *CrI*s | *b* | 99.92% *CrI*s | *b* | 99.92% *CrI*s | *b* | 99.92% *CrI*s | *b* | 99.92% *CrI*s |
| LPMC model | *Promise* vs. *Reply-1* | Speaker | -0.04 | [-0.14,0.07] | 0.01 | [-0.07,0.09] | 0.04 | [-0.03,0.11] | -0.01 | [-0.1,0.07] | -0.01 | [-0.09,0.07] |
| | | Context | 0.02 | [-0.08,0.11] | 0.02 | [-0.06,0.09] | -0.02 | [-0.08,0.05] | 0.01 | [-0.06,0.09] | -0.02 | [-0.09,0.05] |
| MPMC model | | Speaker | -0.02 | [-0.12,0.08] | 0.01 | [-0.08,0.09] | 0.04 | [-0.03,0.11] | -0.01 | [-0.1,0.07] | -0.01 | [-0.09,0.07] |
| | | Context | -0.003 | [-0.09,0.09] | 0.02 | [-0.06,0.1] | -0.02 | [-0.08,0.05] | 0.01 | [-0.06,0.09] | -0.02 | [-0.09,0.06] |
| LPMC model | *Request* vs. *Reply-2* | Addressee | 0.06 | [-0.05,0.16] | 0.01 | [-0.07,0.1] | 0.001 | [-0.07,0.07] | -0.004 | [-0.08,0.08] | -0.03 | [-0.1,0.05] |
| | | Context | 0.04 | [-0.08,0.15] | -0.06 | [-0.14,0.03] | 0.03 | [-0.05,0.01] | 0.04 | [-0.05,0.12] | -0.003 | [-0.08,0.08] |
| MPMC model | | Addressee | 0.05 | [-0.05,0.15] | 0.02 | [-0.06,0.1] | 0.002 | [-0.07,0.07] | -0.004 | [-0.09,0.07] | -0.03 | [-0.11,0.05] |
| | | Context | 0.03 | [-0.07,0.13] | -0.06 | [-0.14,0.03] | 0.03 | [-0.05,0.01] | 0.04 | [-0.05,0.12] | -0.004 | [-0.08,0.08] |

No effect survived the Bonferroni correction (99.92% *CrI*).

**Table S6 | Demographic characteristics of the participants in the lesion study**

| | Lesion laterality | Lesion size (ml) | Chronicity (months) | Age (years) | Sex | Education (years) | MMSE | BDI |
|---|---|---|---|---|---|---|---|---|
| PML1 | Left | 32.15 | 14 | 25 | Female | 16 | 28 | 10 |
| PML2 | Right | 4.48 | 42 | 45 | Male | 15 | 25 | 9 |
| PML3 | Right | 12.82 | 15 | 40 | Female | 9 | 26 | 8 |
| PML4 | Right | 30.03 | 4 | 33 | Female | 16 | 29 | 5 |
| PML5 | Left | 5.26 | 3 | 31 | Male | 16 | 29 | 5 |
| PML6 | Right | 22.33 | 28 | 49 | Female | 16 | 29 | 0 |
| PML7 | Left | 36.46 | 34 | 30 | Female | 16 | 28 | 1 |
| PML8 | Right | 8.54 | 15 | 24 | Male | 16 | 30 | 3 |
| PML9 | Left | 12.8 | 8 | 41 | Female | 9 | 30 | 3 |
| PML10 | Right | 10.58 | 3 | 37 | Male | 16 | 29 | 2 |
| PML11 | Right | 46.55 | 19 | 44 | Female | 16 | 30 | 0 |
| PML12 | Left | 7.15 | 21 | 35 | Female | 16 | 29 | 1 |
| PML13 | Left | 61.58 | 3 | 24 | Female | 15 | 28 | 8 |
| PML14 | Right | 39.66 | 3 | 35 | Male | 15 | 28 | 2 |
| Lesion controls | 7 left/ 6 right | 21.44 (14.22) | 20 (10) | 39 (9) [19, 53] | 6 male/ 7 female | 15 (2) | 29 (1) | 2 (2) |
| Healthy controls | No data | No data | No data | 31 (7) [24, 49] | 9 male/ 18 female | 16 (2) | 29 (1) | 4 (2) |

PML, premotor cortex lesion group; MMSE, mini-mental state examination; BDI, Beck depression inventory. Only participants included in the data analyses are displayed. This table represents individual characteristics of 14 premotor lesion patients and summary characteristics of lesion controls and healthy controls. The numeric characteristics of lesion controls and healthy controls are represented with means, standard deviations (in parentheses), and ranges (in square brackets).
